# Virtual Microbes evolve multiple mechanisms to the same end: anticipating a serial transfer protocol

**DOI:** 10.1101/554766

**Authors:** Bram van Dijk, Jeroen Meijer, Thomas D Cuypers, Paulien Hogeweg

**Keywords:** experimental evolution, serial transfer protocol, eco-evolutionary dynamics, in silico evolution, predicting evolution

## Abstract

**Background:** Experimental evolution of microbes often involves a serial transfer protocol, where microbes are repeatedly diluted by transfer to a fresh medium, starting a new growth cycle. This protocol has revealed that evolution can be remarkably reproducible, where microbes show parallel adaptations both on the level of the phenotype as well as the genotype. However, these studies also reveal a strong potential for divergent evolution, leading to diversity both between and within replicate populations. We here study how *in silico* evolved Virtual Microbe “wild types” (WTs) adapt to a serial transfer protocol to investigate both the generic evolutionary adaptation to such an environment which are independent of prior evolution, and the variety of ways in which the adaptation is implemented at the individual and ecosystem level.

**Results:** We show that all pre-evolved WTs evolve to anticipate the regularity of the serial transfer protocol by adopting a fine-tuned balance of growth and survival. We find that this anticipation can be done in a variety of ways, either by a single lineage or by several lineages in consort. Interestingly, replicate populations of the same WT initially show similar trajectories, but may subsequently diverge along a growth rate versus yield trade-off.

**Conclusions:** We find that all our *in silico* WTs show the same anticipation effects — fitting the periodicity of a serial transfer protocol — but do so by a variety of mechanisms. Our results reveal new insights into the dynamics and relevant selection pressures in experimental evolution, but also highlight how, in an eco-evolutionary context, numerous mechanisms can evolve to the same end.

## Background

In order to see microbial evolution in action, we often rely on experimental evolution under controlled laboratory conditions. The Long-term Evolution Experiment (LTEE)[1] and similar shorter studies[2, 3, 4] have, for example, evolved many generations of microbes using a serial transfer protocol, where microbes are repeatedly diluted and transferred to a fresh medium to start a new growth cycle. Conceptually, if we learn to understand how microbes adapt to such a daily resource cycle, we might one day be able to predict evolution in the lab and — ideally — also in nature. Indeed, a lot of evolution in the lab seems remarkably reproducible, where microbes show parallel adaptations both on the level of the phenotype as well as the genotype [5, 6, 7, 8, 9, 10, 4, 11]. However, there also seems to be strong potential for divergent evolution, leading to diversity both between and within replicate populations [12, 13, 14]. Diversification events within populations often entail cross-feeding interactions [15, 16, 12, 17, 13, 18], where species emerge that grow on metabolic by-products. These cross-feeding interactions are increasingly well understood with the help of metabolic modeling and digital evolution [19, 20]. A recent metagenomicstudy has revealed even more coexisting lineages in the LTEE than were previously reported [21]. It is however not yet clear whether all these polymorphisms are the result of uni-directional cross-feeding interactions, or if other mechanisms could drive coexistence in a simple experiment such as a serial transfer protocol. Prior to being subjected to lab conditions, the microbes used in the aforementioned experimental studies have all had a long evolutionary history in natural environments, experiencing harshly fluctuating and — more often than not — unfavorable conditions. While a serial transfer protocol such as that of the LTEE at a first glance selects mostly for higher growth rates when resources are abundant (i.e. during the log phase), there is also selection to survive when resources are depleted and the population no longer grows (*i.e.* during the stationary phase). In fact, given the unpredictable conditions found in nature, some of the ancestors of *Escherichia coli* might have survived precisely because they diverted resources *away* from growth. Indeed, *E. coli* does exactly this during the stationary phase by means of the stringent response, regulating up to one third of all genes during starvation [22]. This response lowers the growth rate, but promotes efficiency and survival (*i.e.* a higher yield). While most microbes have ways to deal with starvation, the physiology of growth arrest varies a lot across different microbes (for an excellent review, see [23]). Responses to starvation, as well as other features that organisms have acquired during their evolutionary history (such as gene clustering, gene regulatory network architecture, metabolic regulation), might strongly influence the adaptation and reproducibility we observe in the lab today.

What do we expect when a complex, “pre-evolved” microbe adapts to a serial transfer protocol like the LTEE? We here use Virtual Microbes in order to firstly mimic natural evolution, acquiring Virtual “wild types” (WTs), which we then expose to a serial transfer protocol (see methods). We do so in order to obtain a fresh perspective on *what* is being selected for, *how* this target can be achieved, and which *generic features* might appear in spite of evolutionary contingencies. We find that the evolved WTs — which are both genotypically and phenotypicall diverse — also lead to a variety of different solutions when subjected to a serial transfer protocol. More specifically, we see many alternative paths in terms of a growth dynamic trajectories, speciation and regulation. Despite this diversity however, all WTs evolve the same anticipation towards the serial transfer protocol, timing their growth rate, yield, and survival to accurately fit the daily cycle. Whereas some WTs do this by means of clever gene regulation, others diverge into specialised growth and survival strains, and other simply time their resource consumption as to not over-exploit the medium. In short, our WTs all recognized and exploited the regularity of the serial transfer protocol, anticipating resources to be available as usual, but they solve this challenge by a variety of different mechanisms.

## Results

### Using Virtual Microbes to search for generic patterns

In this study we use Virtual Microbes, a model of the eco-evolutionary dynamics of microbes (see methods). In short, the Virtual Microbe model is unsupervised, meaning that it aims to combine relevant biological structures (genes, genomes, metabolism, mutations, ecology, etc.) without a preconceived notion of “fitness”, which is instead an emergent phenomenon. By not explicitly defining what the model *should* do, it allows for a serendipitous approach to study microbial evolution. Our main objective in this study is to elucidate *generic patterns* of evolution in a serial transfer protocol, and to investigate the extend to which these are constrained by prior evolution. In order not to lose track of the objective of *finding generic patterns*, we refrain from discussing and analysing every mechanistic detail, and instead focus on major observables and discuss some illustrative cases. Before we start evolving Virtual Microbes in a serial transfer protocol, we first evolved a set of Virtual “wild types” (WTs). Instead of optimizing these WTs solely for high growth rates, we here mimic natural circumstances by fluctuating resource conditions (Figure 2A). When too little resource is available, the Virtual Microbes cannot grow. When too much resource is available however, the Virtual Microbes run the risk of accumulating too high concentrations of metabolites, resulting in increased death rates due to toxicity. To avoid extinction, we divided the total grid into four sub-grids. In these sub-grids, the two resource metabolites A and C (yellow and blue in Figure 1A) change in their influx rates with probability 0.01. Both these resources can be converted into building blocks (purple) required for growth. Maximally flourishing Virtual Microbes live on average 100 time steps. Thus, a healthy Virtual Microbe experiences on average one fluctuation in resource conditions in its lifetime (see full configuration in S1). As the rates of influx span four orders of magnitude, conditions will vary from very favourable to very poor. This in turn depends on which resources the evolved Virtual Microbes like to consume (and at which rate), whether or not there is too much or too little resource, and whether or not space for reproduction is available. All in all, this results in an unsupervised evolutionary process where there is no prior expectation of what metabolic strategy or gene regulatory networks might be best suited to survive. Because competitive fitness is not a priori defined, we can study what will be the long-term target of the eco-evolutionary dynamics, not in terms of fitness, but in terms of what the Virtual Microbes evolve *to do*.

**Figure 1.**
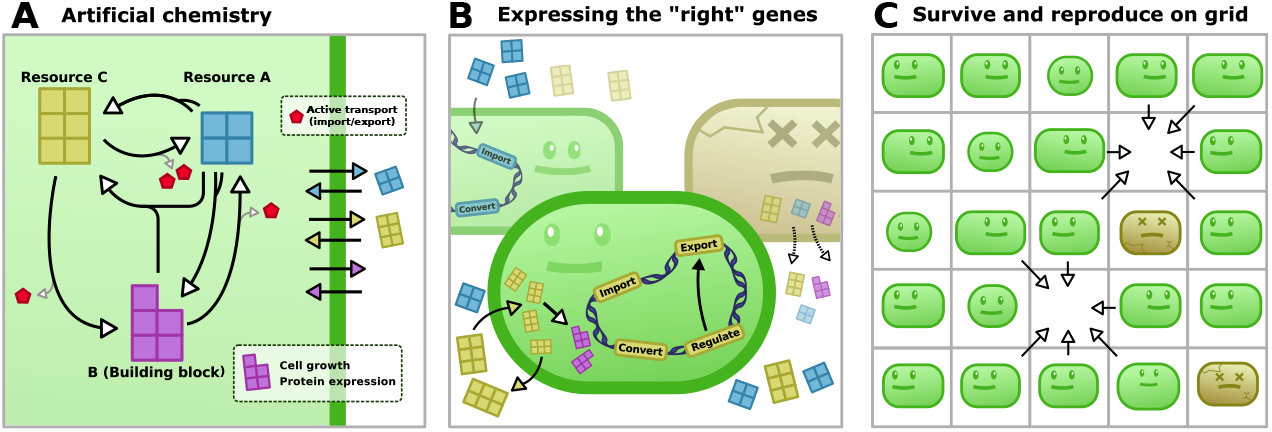
Virtual Microbes model overview. **A)** At the basis of the Virtual Microbe model is an artificial “metabolic universe”, describing all the possible reactions that can be catalysed. Resources (yellow and blue) are fluxed in, but building blocks (purple) and energy (red) must be synthesized to express proteins and transport metabolites across the membrane, respectively. **B)** A Virtual Microbe only needs to express a subset of all possible reactions to be viable, and that no metabolic strategy is necessarily the “right” one. **C)** The individuals grow and reproduce on a spatial grid, and can only reproduce when there is an empty spot. Death happens stochastically or when a cell has accumulated toxicity by having excessively high concentrations of metabolites. Since only cells that have grown sufficiently are allowed to reproduce, we simulate evolution with no prior expectation.

Using this protocol, we evolved the same initial clone in the exact same “random” resource fluctuations, *only* varying the mutations that happened across ~10.000 generations of evolution. This produced 16 distinct WTs with their own evolutionary history, which we then expose to the serial transfer protocol (Figure 2B). Despite experiencing precisely the same fluctuations, no two WTs evolved to be the same. For example, we observe a great diversity in gene content, kinetic parameters of enzymes, gene regulatory networks and their complexity, and responses to environmental stimuli. The core metabolism is however strikingly similar across WTs, always consisting of a simple metabolic cycle. The rates of building block production and death rates are also very similar across all WTs (Figure S3). In other words, it appears that there are many different ways to be fit, and that no solution is evidently better. The similarities and differences between our WTs are summarized in Figure 2C, but we discuss this in more detail in Supplementary Section 1.

**Figure 2.**
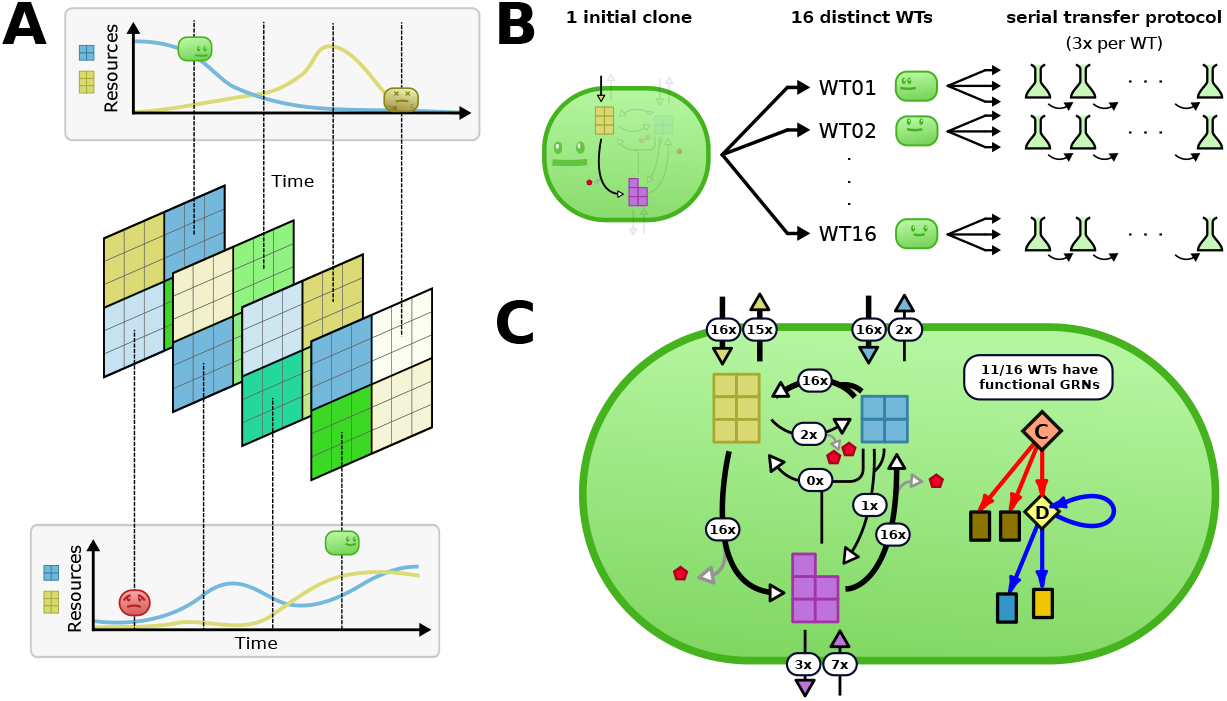
Evolution of Virtual “wild types” under naturally unpredictable and fluctuating resource conditions. **A)** Natural evolution is mimicked by (harsly) fluctuating resource conditions, resulting in a wide variety of resource conditions. The (actual) grid is 40×40, with four 20×20 subspaces where the rates of influx vary stochastically. These subspaces do not impede diffusion of metabolites or reproduction. The fluctuations of the A and C resource (blue and yellow respectively) are independent, resulting in a variety of different conditions. Building blocks (purple) must be synthesized for growth and protein expression. **B)** We repeat the evolution in natural conditions 16 times starting from the same (minimally viable) initial clone (varying the mutations that happen) yielding 16 distinct WTs. These WTs are later transfered to a serial transfer protocol. **C)** In the white labels we show how many of the evolved WTs adapted to use particular reactions. The thicker arrows represent the shared core genome which consists of two resource importers, a metabolic cycle, and a C-exporter. Transcription factors (diamonds) were always present across WTs, but only 11/16 WTs visibly display changes in gene expression correlated with changes in the environment.

### Long-term evolution experiment *in silico*

After evolving a variety of different WTs, we mimic a serial transfer protocol like that of the LTEE. With regular intervals, all but 10 percent of the cells are removed, while at the same time refreshing the medium. Although time in Virtual Microbes has arbitrary units, we will refer to this process as the “daily” cycle from this point forward. Early in the day, during the log phase, high growth rates are very rewarding as there is a lot of opportunity to reproduce. However, once the population has reached stationary phase (having consumed all resources), it is favourable to survive and to not invest in growth any further. We will focus on how our WTs adapt to these alternating selection pressures. The results discussed here are found for a variety of different medium conditions (*e.g.* also see Table S2). In the main text however, we present the 50 time step serial transfer protocol where the medium contained both resources (A and C), as this was a condition on which all WTs could be cultivated, ensuring equal treatment. We focus on the generic features of the adaptation towards this protocol, and how specific WTs and contingent factors from their evolutionary history shape these outcomes.

### All wild types evolve to anticipate the serial transfer protocol

After 800 days of evolving in a serial transfer protocol, we compare the ancestral WTs with the evolved populations. We firstly look at some of the well-known growth dynamics of microbes: the lag-, log-, and stationary phase (Figure 3A). As most experimental evoultionary studies in the lab, we too observe a decreased lag phase and an increased growth rate. The increased growth rate in the evolved population results in an earlier onset of the stationary phase, which therefore takes much longer than for their WT ancestors. Eventually, this leads to a phase where the cell count decreases again (death phase), revealing a decrease in survival for the evolved populations. To further study how this decreased survival comes about, we next investigated the dynamics of average cell volumes, which are indicative of the “health” of the population: cell volume determines the ability to divide (minimal division volume) and survive (minimal viable volume). A first interesting observation is an increase in average cell volume during the log phase (Figure 3B-C), which is also one of the first results from the LTEE[24]. However, after this increase in cell volumes during the log phase, evolved populations display a clear decrease in cell volumes, either at the end of the day (Figure 3B), or during the whole stationary phase (Figure 3C). Indeed, if we expose the populations to prolonged starvation by extending the day, the evolved populations die shortly after the anticipated serial transfer, while their WT ancestors survived for much longer (Figure 3B-C, right-hand side). Interestingly, we observed that the cell volume at the time of transfering the cells to a fresh medium (henceforth ‘transfer volume’) fall into two distinct categories. In the high yield scenario (Figure 3B), cell volumes are maintained *above the division volume* until the very end of the day, whereas the low yield scenario leads to a transfer volume that is *just above minimal.* While the distribution of these observed transfer volumes across ancestral WTs are mostly high (Figure 3D, left-hand side), the evolved cells show a bimodal distribution (Figure 3D, right-hand side). The WTs evolved to either be ready to immediately divide at transfer (Figure 3B), or exploit as much resource as possible while remaining above the minimal viable volume (Figure 3C). Despite this difference, both alternative strategies have evolved to anticipate the regularity of the serial transfer protocol. Indeed, when the extended yield (the total biomass that was generated after prolonged starvation) is measured, it shows a consistent decrease across all evolved populations (Figure 3E) relative to the WTs, as these long term effects are now masked from natural selection. We found that this anticipation effect did not depend on details in the protocol, such as the length of the daily cycle or the number of resources used (Figure S5, Table S2). This reveals that a key selection pressure in a serial transfer protocol is not only growth as fast as possible, but also remaining viable until the next day, anticipating the next food supply.

**Figure 3.**
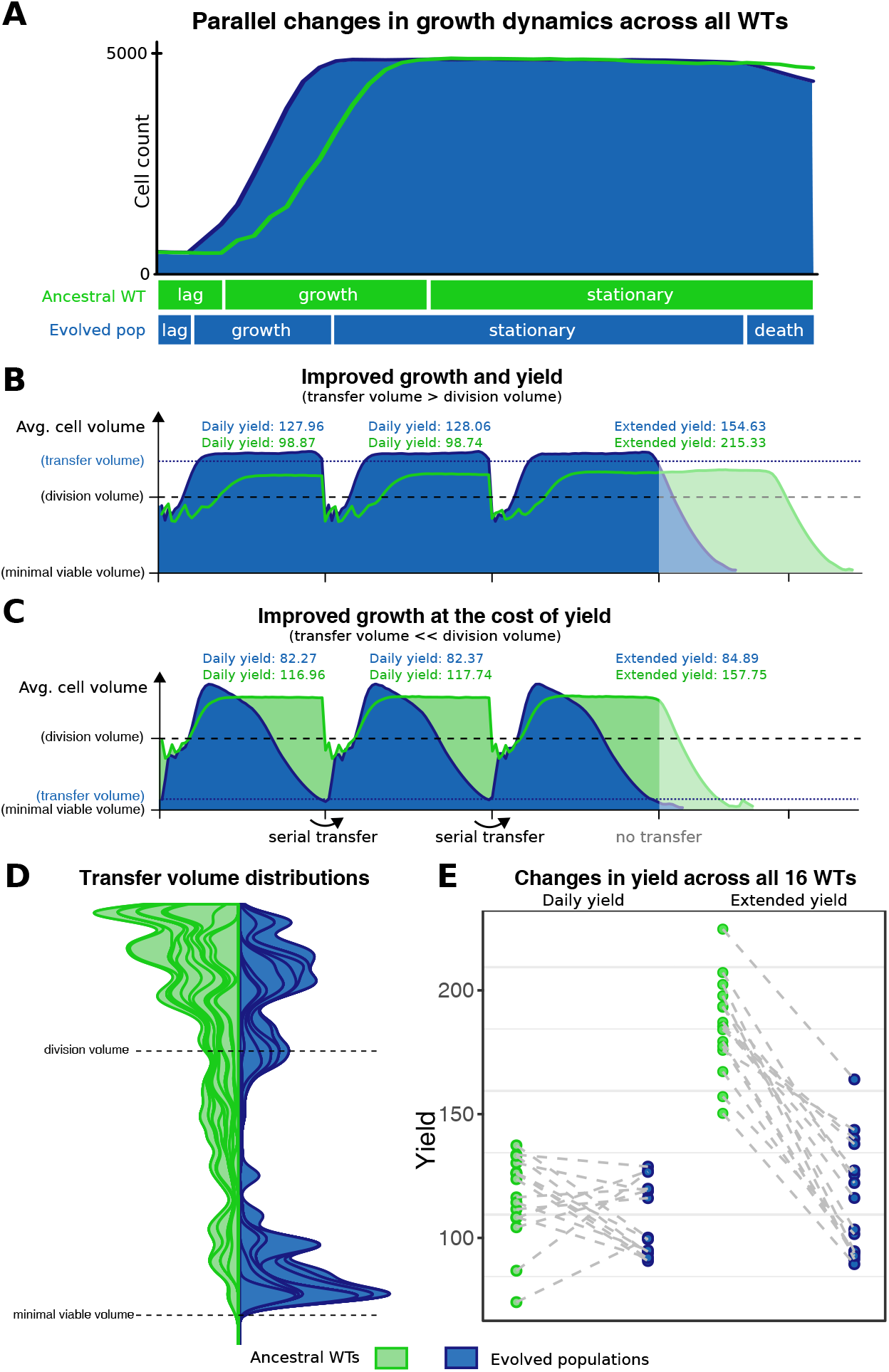
Virtual Microbes adapt to anticipate the regularity of a serial transfer protocol. **A)** Growth dynamics of WT (green, day 10) and evolved populations (blue, day 760) in terms of cell counts. (WT03#1 taken as an illustrative example). **B-C)** Two WTs (green) and the population after prolonged evolution in the serial transfer protocol (blue) are shown as an illustration of the anticipation effects. Over the course of 3 cycles, the average cell volume is plotted against time for the ancestral WT (green) and for the evolved population (blue). The y-axis (cell volume) indicates the minimal viable volume and division volume (which are fixed for the model), and the evolved transfer volume (as measured at the end of the third cycle). Daily and extended yield are measured as defined in the method section. After the third cycle, serial transfer is stopped (transparent area), showing decreased survival of the evolved populations with respect to their ancestor. **D)** Stacked density distributions are plotted for the transfer volumes both early (transfer 0-40, green) and late (transfer 760-800, blue). **E)** The evolved changes in yield both “daily” (within one cycle of the protocol) and “extended” (after prolonged starvation) for all 16 WTs.

### WTs have distinct trajectories toward a growth-yield trade-off

The two extreme categories of cell volume dynamics from Figure 3 suggest a tradeoff between growth and yield. We next investigate how our different WTs evolve towards this trade-off, and how reproducible these trajectories are. For this, we repeated the serial transfer protocol 3 times for each WT, and follow the trajectories over time. After ~800 serial transfers, all populations have adapted along a tradeoff between growth and yield (Figure 4A, p << 10e-16, R^2^ = 0.54). This trade-off was not observed during the first cycle of the protocol, which instead shows a positive correlation between growth and yield (Figure 4B, p << 10e-5, R^2^ = 0.32). Most WTs predictably evolve towards the trade-off by improving both growth and yield (*e.g.* by importing more resources, or producing more building blocks), but subsequent evolution is very WT-specific. Many WTs maintain high yield (*e.g.* Figure 4C), but others consistently trade off yield for a higher growth rate (Figure 4D). For instance, importing even more resources can improve growth even further, but leads to prolonged starvation and/or toxicity. Lastly, some WTs are showing variable trajectories after having arrived at the trade-off (Figure 4E, Figure S6). Taken together, these results illustrate how prior adaptations strongly shape the way subsequent evolution plays out. Evidently, specific WTs more readily give rise to certain solutions, having specific adaptations in their “mutational neighbourhood”. This is also illustrated by two WTs that repeatedly gave rise to mutants with extremely high, but unsustainable growth rates, causing multiple populations to go extinct (black crosses in Figure 4). In summary, some WTs adapt predictably to the serial transfer protocol, while others have diverging evolutionary trajectories and can reach different solutions. The consistency of WTs in combination with the diversity of trajectories illustrates how prior evolution can bias — but not necessarily constrain — subsequent evolution.

**Figure 4.**
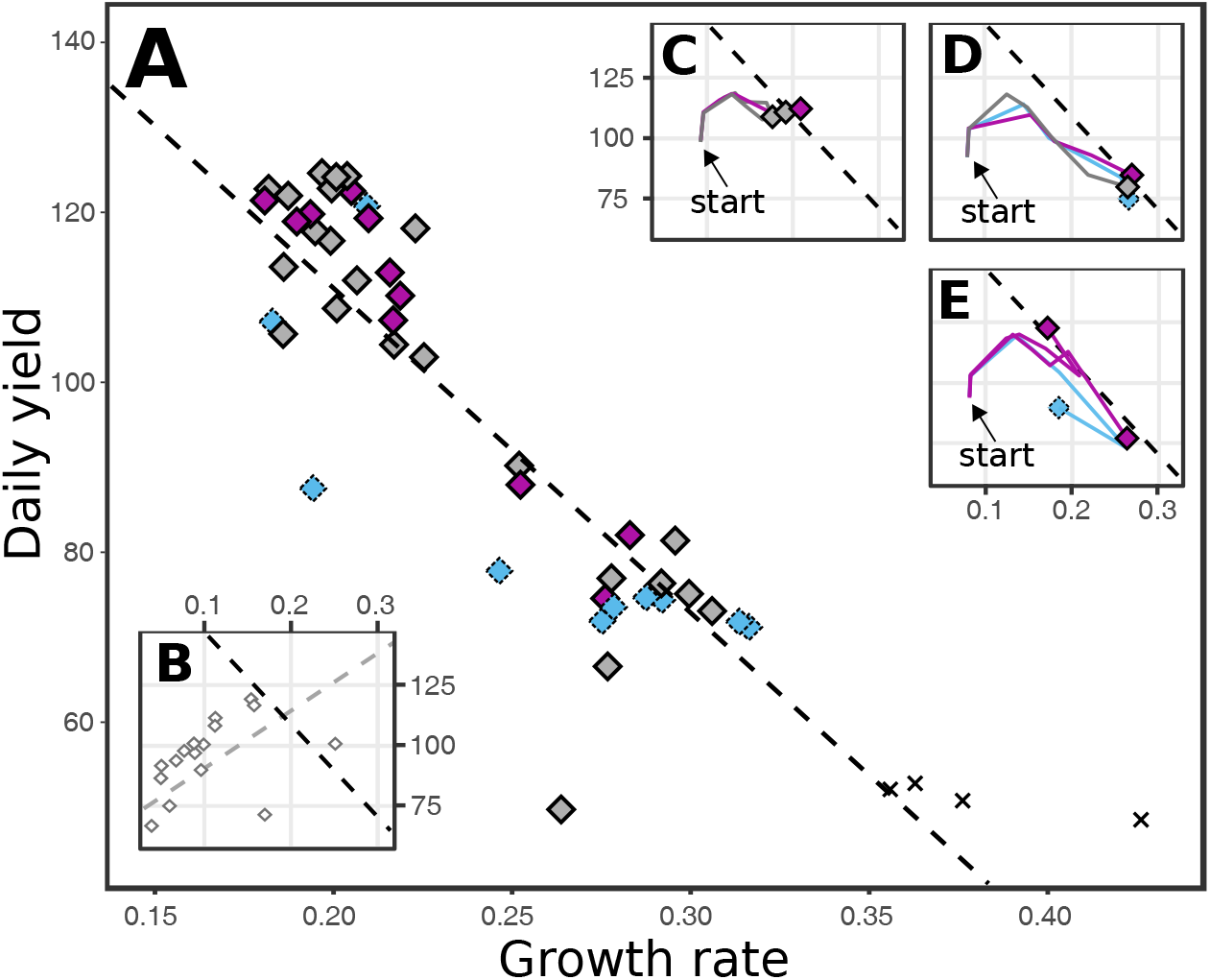
Prior evolution determines trajectories of adaptation towards a serial transfer protocol. **A)** Growth rate (average building block production during log phase) is plotted against daily yield (average population biomass within a single cycle), for all the 48 experiments after adaptation to 800 serial transfers. The black dotted line is a linear regression model (p << 10e-16, R^2^ = 0.54). **B)** Shows the initial points for all 16 WTs, which actually have a positive correlation between growth and yield (p << 10e-5, R^2^ = 0.32) instead of the negative correlation (black dotted line). **C-E)** These insets display how the repeated evolution of most WTs produces similar trajectories towards the trade-off (time points are day 0, 20, 40, 100, 200 and 800), ending in either high daily yield (C) or low daily yield (D). A minority of WTs diverge after reaching the trade-off, and thus show more diverse trajectories when repeated (E). The colours of the end point symbols depict different modes of adaptation as discussed in the next paragraph (grey = no coexistence, blue = quasi-stable coexistence, purple = stable balanced polymorphisms, black cross = extinction).

### Polymorphism based anticipation by evolution growth and survival strains

So far we have only looked at population averages. Next, we study the dynamics of lineages and the evolved dynamics within cells. To track lineages we tag each individual in the population with a neutral lineage marker at the start of the experiment (analogous to DNA barcoding). When a single lineage reaches fixation, we reapply these neutral markers, allowing us to quickly detect long-term coexistence. Moreover, these neutral markers allow us to study which arising mutants are adaptive in the different phases of the growth cycle. In Figure 5A we show dynamics of neutral lineage markers that are frequently redistributed when one lineages fixates in the population, indicating that there is no long-term coexistence of strains. In contrast, figure 5B displays a repeatedly observed quasi-stable coexistence, where two lineages coexist for some time, but coexistence was not stable in the long-term. Lastly, Figure 5C shows stable, long-term coexistence, where two lineages coexisted until the end of the experiment. Coexistence (either quasi-stable or stable) was observed in 21 out of 44 extant populations (Figure 5D).

**Figure 5.**
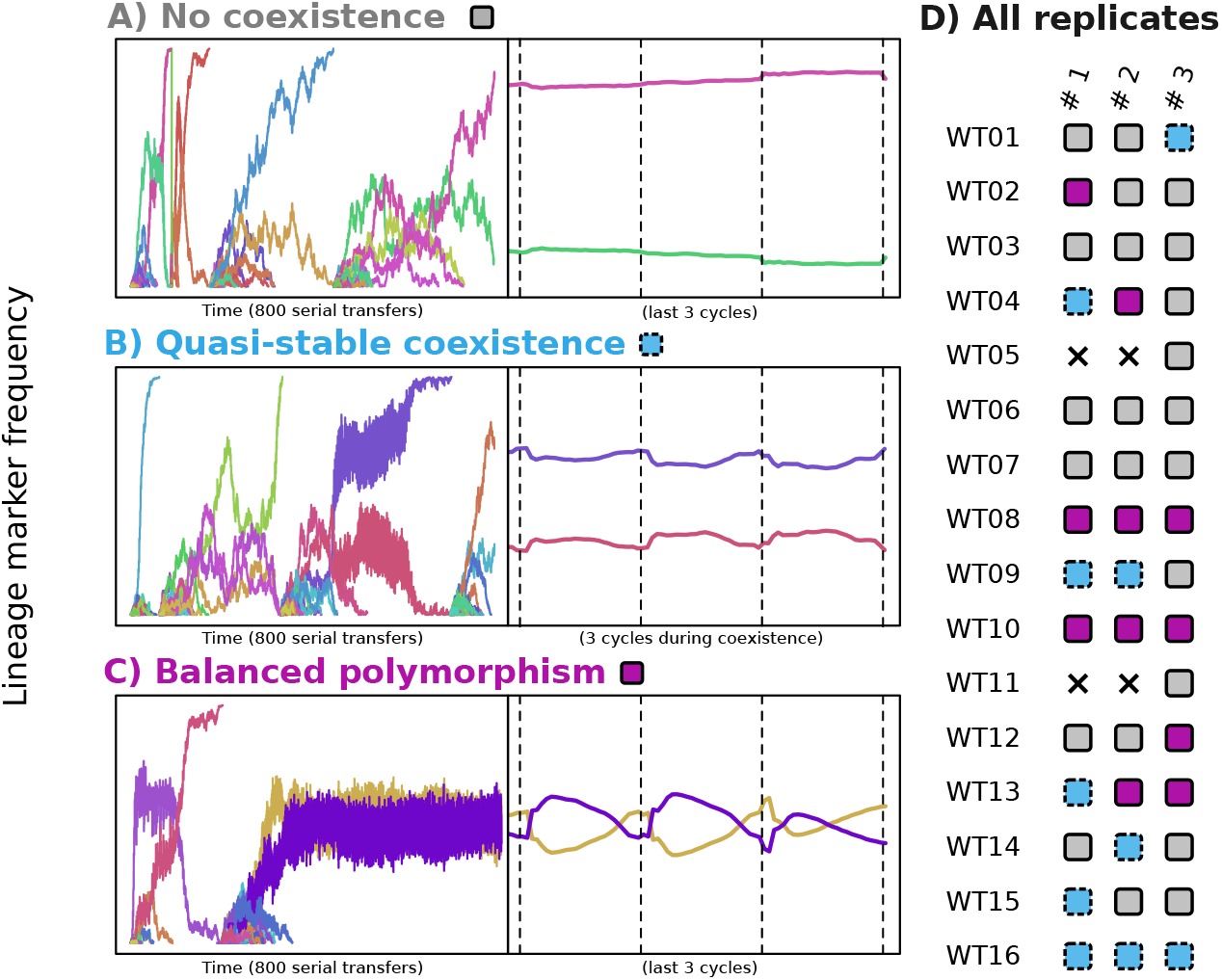
Dynamics of neutral lineage markers reveal balanced polymorphisms based on the daily cycle. **A-C)** Neutral lineage markers (random colours) frequencies are plotted along 800 serial transfers (left hand side) and along 3 cycles. Panel A shows an example with no coexistence which is found in 23 out of 44 replicates, and panel B and C show (quasi-)stable coexistence, found in the remaining 21 replicates. **D)** shows these 3 possible outcomes for all 3 replicates of 16 WTs (grey = no coexistence, blue = quasi-stable coexistence, purple = stable balanced polymorphisms, black cross = extinction) Four replicates went extinct during the serial transfer experiment due to over-exploiting of the medium (black crosses).

By zooming in on the dynamics of coexisting lineage markers over a shorter time span (Figure 5B-C, right-hand side), we can better understand how these lineages stably coexist. Notably, one lineage is dominating during log phase, while the other lineage performs better during stationary phase. In other words, the lineages have specialized on their own temporal niche. We find that these dynamics can be the result of three mechanisms (or combinations thereof): 1) cross-feeding on building blocks, 2) specialisation on resource A or C, 3) based on the growth vs. yield tradeoff. Cross-feeding dynamics always resulted in quasi-stable coexistence (such as depicted in 5B), and never resulted in the balanced polymorphism as depicted in Figure 5C), while the other two mechanisms (resource specialisation and growth vs. yield differentiation) most often resulted in long-term coexistence where the lineages perform better together than they do alone (Figure S8). While specialisation on different resources is a well known mechanism for negative frequency dependent selection, it is far less evident how a growth vs. yield trade-off would result in a fully balanced polymorphism. Mutants with higher growth rates but elevated death rates have a very distinct signature of increasing in frequency early in the daily cycle and decreasing to much lower frequencies during the stationary phase (Figure S7A), as apposed to lineages that increase in frequency throughout all phases of the cycle (Figure S7B). While such mutants readily arise across our experiments, they often have difficulty rising to fixation due to an increasing duration of the stationary phase. In the meantime, a slower growing lineage with lower death rates can be optimized to utilize resources at low concentrations during stationary phase. Evidently, these dynamics can give rise to a balanced polymorphism that does not depend on resource specialisation, as it is also observed in our experiments with a single resource (Table S2). Indeed, Figure 5A illustrates how two lineages with more than a three-fold different death rates can stably coexist.

Besides this speciation on the basis of the growth vs. survival trade-off, we also found well-known mechanisms of speciation, such as cross-feeding [11, 16, 18], canabalism[13], or other resources in the medium [15, 25]. The nature of the coexistence can differ strongly across WTs and replicated experiments. For example, since *de novo* gene discoveries were disabled during this experiment, cross-feeding on building blocks is only possible if the ancestral WT had the necessary importer for said building block, which was true only for 7/16 WTs. Similarly, even though all WTs have the necessary importers for both the A and C resource, only one WT consistently diverged into an A- and C-specialist (WT10). While other WTs have multiple gene copies for these importers, WT10 had only 1 copy of both genes, making the loss-of-function mutations readily accessible. In conclusion, all polymorphic population anticipate the serial transfer protocol, but do so by a variety of mechanisms. However, they all have in common a *generic pattern* of strains which time growth and survival strategies in relation to each other to precisely finish the available food resources by the end of the day.

### Single lineage anticipation by tuning and trimming the gene regulatory network

The previous section illustrates how multiple lineages can coexist because the predictable serial transfer protocol produces temporal niches. However, many of our WTs do not show any tendency to speciate like this, and instead always adapt to the serial transfer protocol as a single lineage (Figure 6D). In order to better understand this, we will now look at the intracellular dynamics of WT07, and how it changes when adapting to the protocol. WT07 is one of the more “clever” WTs with a relatively complex GRN, and displays strong responses in gene expression when exposed to “natural” fluctuations. In Figure 6B we show that WT07 consistently adapts to the protocol by switching between two modes of metabolism, where importer proteins are primed and ready at the beginning of the cycle, and exporter proteins and anabolic enzymes are suppressed during stationary phase. Despite some differences in the evolved GRNs, the evolved protein allocation patterns are virtually indistinguishable across the three replicates. Interestingly, although no parallel changes were observed in the kinetic parameters of proteins, we do observe the parallel loss of a energy-sensing transcription factor as well as increased sensitivity of the TF that senses the external resource C. In other words, evolution apparently happened mostly through loss, and tuning and trimming of the GRN. Modulation between two metabolic modes allows this single lineage to switch between log and stationary phase, occupying both temporal niches. Indeed, a second lineage never appeared for this WT (Figure 6B and Table S2).

**Figure 6.**
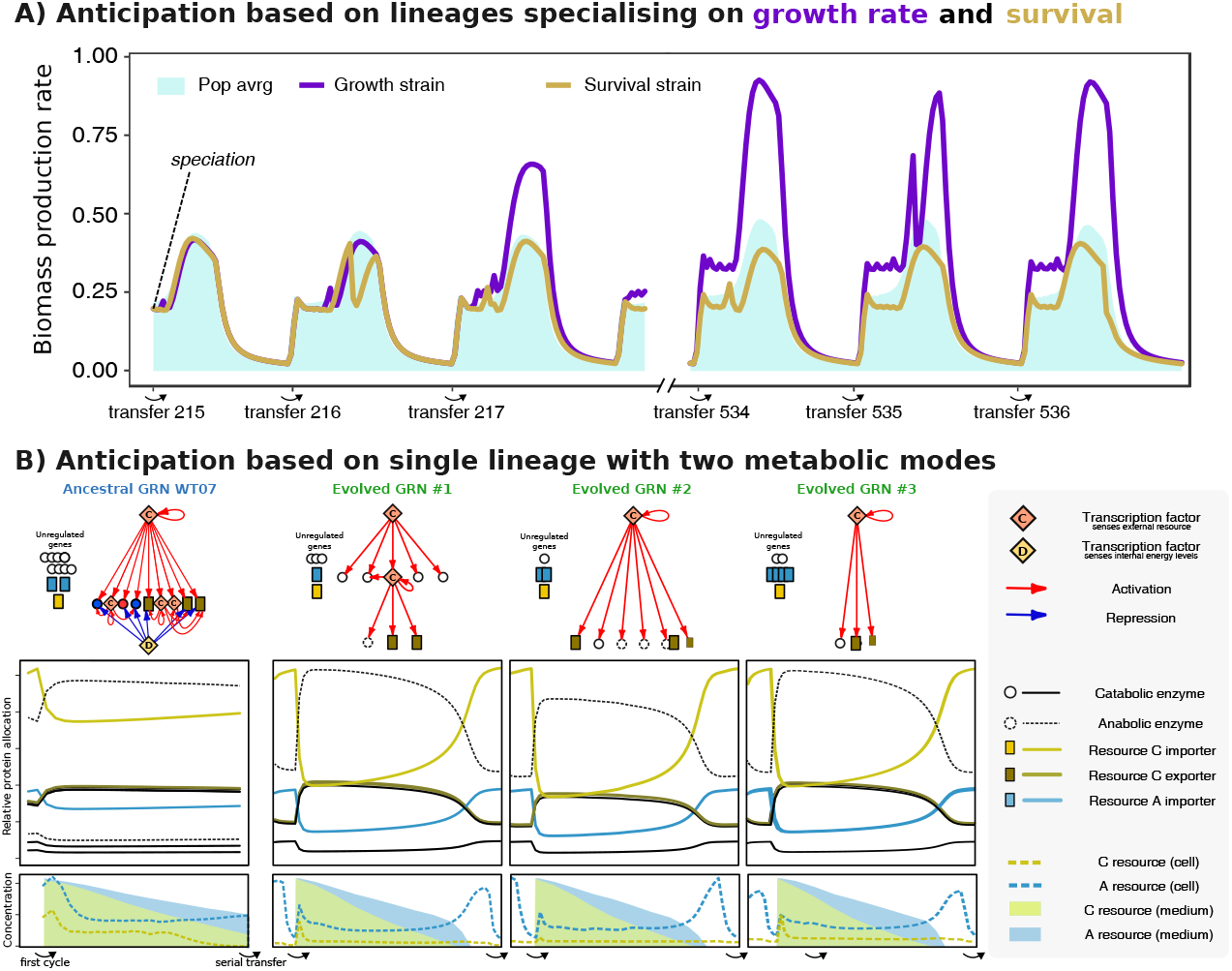
Anticipation can entail polymorphism or a single lineage that switches between two metabolic modes. **A)** Two lineages occupy different niches on the growth vs. yield trade-off WT02#01 diverges into a slow growing lineage (yellow lineage, average death rate ±0.015) and a faster growing lineage with elevated death rates (blue lineages, average death rate ±0.048), together anticipating the serial transfer protocol. **B)** A single-lineage anticipates the daily cycle by trimming and tuning the gene regulatory network. All three replicates of WT07 anticipate as a single lineage with two metabolic modes. The ancestral gene regulatory network (GRN), protein allocation dynamics, and resource concentrations are displayed for WT07 during the first cycle of the serial transfer protocol, and for the three replicated experiments after 400 serial transfers.

Strikingly, a GRN does not necessarily lead to a single lineage adaptation. For example, another regulating wild type (WT13) repeatedly evolved into multiple coexisting lineages, while maintaining the ability to regulate gene expression. *Vice versa*, non-regulating wild types (WT01 and WT15) also evolved single-lineage anticipation. Hence, even though the GRN of WT07 has a major impact on the repeatability of single-lineage adaptation (as illustrated in Figure 6B), the presence of a functional GRN is neither sufficient nor necessary for single lineage adaptation.

## Discussion

In this study we have taken a serendipitous approach to study how microbes adapt to a serial transfer protocol, and to what extent this is determined by their evolutionary history. The Virtual Microbe modelling framework serves this goal by not explicitly defining the concept of fitness. Instead, it builds up biology from the bottom up by implementing basic biological features and their interactions. We observe that regardless of their evolutionary history, all WTs learn to anticipate the regularity of the serial transfer protocol by evolving a fine-tuned balance between high growth rate and yield. Long-term survival without food, which is now masked from natural selection, always deteriorates after prolonged exposure to such a protocol. We next show that, if the same WT is repeatedly evolved in a serial transfer protocol, it has similar trajectories towards a growth versus yield tradeoff, but may subsequently diverge along it. Polymorphisms within populations are frequently observed, which can happen by means of cross-feeding interactions, resource specialisation, or growth vs. yield specialisation. We furthermore find that coexisting lineages are dependent on each other, as they would perform better in the presence of the other. In general, our results are robust to details in the serial transfer protocol, such as using only a single resource, or varying the interval between transfers (see Table S2). The anticipation effects therefore appear to be generic features of microbes exposed to prolonged evolution in a serial transfer protocol. Moreover, the concept of microbial populations anticipating predictable changes has also been observed in previous *in silico*[26] and experimental studies[27]. Combined with diversification and bet hedging strategies, anticipation might well play an important role in natural populations, the details of which are yet to be elucidated[28].

How do our results map onto experimental evolution in the lab? *E. coli* Bc251 has been subjected to a daily serial transfer protocol for over 30 years (~70.000 generations) in the LTEE. Many of our observations are remarkably similar to the LTEE, such as the improved growth rate and cell sizes during the log phase[24], the (quasi-)stable dynamics of coexisting lineages[21], and “leapfrogging” dynamics (*e.g.* Figure 5A-B) where an abundant lineage is overtaken by another lineage before rising to fixation[29, 30]. The comparison with respect to the growth versus yield dynamics and the anticipation effects discussed in this work is however less straightforward. We have observed how all our WTs quickly evolve to be maximally efficient given our artificial chemistry, and only subsequently diverge along the apparent growth versus yield trade-off (see Figure S6). For this strain of *E. coli*, growth and yield have continued to improve so far, and although a trade-off has been observed *within* the populations[31], no growth versus yield trade-off between the replicate populations has been observed yet. Likewise, our interesting results on the evolution of anticipation could not be corroborated as of yet (T. Hindre and D. Schneider, personal communication, November 2018). Nevertheless, we propose that anticipation of periodic environmental change, and a growth versus yield trade-off, provides testable hypotheses for the future of the LTEE, and similar experimental studies.

Cross-feeding interactions are commonly observed in the LTEE and similar studies [18, 11, 11, 17], and modeling has shown that this adaptive diversification involves character displacement and strong niche construction[19], and can furthermore strongly depend on the regularity of a serial transfer protocol [20]. While similar cross-feeding interactions are observed in some of our *in silico* experiments, we also found balanced polymorphisms involving one lineage with high growth rates during log phase and a slower growing lineage which performs better in stationary phase. This can happen by means of resource specialisation, or purely on the basis of a growth versus yield specialisation which does not require cross-feeding or cannibalism. While the resource specialisation is only relevant to experimental studies that use more than one carbon source, the growth versus yield diversification also happens on a single resource (Table S2). Indeed, other studies have also suggested the importance of these dynamics, such as the coexistence of respiratory and fermenting strains in *Saccharomyces cerevisiae* [32] in a chemostat, and the presence of multiple selection pressures in a mathematical model of a serial transfer protocol [33]. Our findings show that these dynamics can emerge in a more complex eco-evolutionary setting. It however remains to be seen if such diversification happens in experiments such as the LTEE. Earlier work on the LTEE has shown that, although no significant negative correlation was found for an evolutionary trade-off of growth and yield, isolated clones from *within* a population do display a negative correlation [31], suggestive for the dynamics we observed in Virtual Microbes. With a great deal of polymorphisms in the LTEE left unexplained so far[21], we thus suggest this as a search image for future evolutionary experiments.

Much to our surprise, we failed to observe any consistent difference between the WTs that had evolved regulated gene expression before the onset of the serial transfer protocol, and those that did not. Even though we observed that gene regulation was often tuned to anticipate the serial transfer protocol (Figure 6), this solution did not appear to be any more productive than solutions that used no gene regulatory mechanisms. Because all the kinetic parameters of enzymes (K_*m*_, V_*max*_, etc.) in the Virtual Microbes are freely evolvable, it is possible that metabolic regulation of homeostasis plays a very important role in Virtual Microbes, and can in many ways be just as “good” as a functional gene regulatory network. Furthermore, we noticed that for certain WTs, a change in metabolism could bypass protein expression by means of kinetic neofunctionalistaion of paralogous genes. Although such a solution does waste more building blocks on the continuous production of protein, it is also much more responsive to environmental changes. Indeed, recent work has shown that adding enzyme kinetics to models of metabolic fluxes leads to much more robustness without leading to a loss in adaptive degrees of freedom [34]. However, the LTEE has revealed many changes in the GRN [7] and global transcription profiles. Thus, the GRN appears to add knobs and buttons for evolution to push [9], but it does not change what is actually being selected for: anticipating the serial transfer protocol.

### Moving towards an eco-evolutionary understanding

The dynamics of Virtual Microbes expose that even a simple serial transfer protocol entails much more than sequentially evolving faster and faster growth rates. Instead, adaptation is an eco-evolutionary process that strongly depends on prior evolution, timescales, the presence of other competitors and mutants, and transient fitness effects. In accordance with these dynamics, temporal positive selection for certain alleles can be inferred from a metagenomics study on the LTEE[21]. Strikingly, although competition experiments favoured the evolved population over the ancestral WTs in almost all cases, there were exceptions to this rule. It is therefor possible that the ancestral WTs perform better in such an experiment, but that this does not describe a stable eco-evolutionary attractor. Indeed, survival of the fittest is an eco-evolutionary process where a lineage interact with other lineages, or with mutants, through changes in the environment. In the LTEE, faster growth might become less and less important as the years pass, perhaps making the aforementioned interactions between lineages increasingly relevant. Other recent studies have recently elucidated the importance of eco-evolutionary dynamics[35], and how this can readily give rise to coexistence of multiple strains which could not have formed from a classical adaptive dynamics perspective[36, 37]. Indeed, metagenomics have revealed much more diversity in the LTEE than previously anticipated[21]. Shifting focus from competition experiments towards the ever-changing selection pressures that emerge from the eco-evolutionary dynamics and interactions, will make the field of experimental evolution harder, but more intriguing, to study.

## Conclusions

We have studied how *in silico* WTs of Virtual Microbes adapt to a serial transfer protocol like that of the LTEE. While the LTEE has shown a sustained increase in competitive fitness, and intensive research displays how the evolved clones are still improving with respect to their ancestor up until today [38, 39, 40]. It is however still unclear what the actual selection pressures are which are at stake. Our experiments have generated a novel hypothesis that what is being selected for is not necessarily high, but balanced growth, which happens with a great variety of underlying mechanisms. A fine-tuned balance between growth and yield causes an accurate anticipation of the serial transfer protocol, which happen by “clever” single lineages, or by multiple lineages that arise on the basis of the growth versus yield trade-off. Taken together, our results reveal important insights into the dynamics and relevant selection pressures in experimental evolution, advancing our understanding of the eco-evolutionary dynamics of microbes.

## List of abbreviations

LTEE: : Long Term Evolution Experiment (first published by R Lenski, 1991)
WT: : wild type (plural: WTs)
TF: : Transcription Factor (plural: TFs)
GRN: : Gene Regulatory Network (plural: GRNs)

## Methods

A full description of the model and underlying equations is available online (bitbucket.org/thocu/virtual-microbes and https://virtualmicrobes.readthedocs.io). Here we summarize the sections of these documents that are relevant to this study.

### Finding generic patterns of evolution

Experimental evolution is, of course, done on organisms that have evolved for a long time under a wide variety of conditions. These studied organisms all have their own evolutionary history, and differences in how they deal with starvation, stress, changes in resource etc. With Virtual Microbes we are able to evolve a *de novo* set of “wild types” (WTs), adapted to live in such severely fluctuating resource conditions. We can then explore how these WTs adapt to experimental evolution, and find generic patterns of evolution. To find generic patterns without being biased towards specific solutions, the biology of Virtual Microbes build-up from many levels with many degrees of freedom. One downside of this strategy can be that every simulation (like biological evolution itself) results in a slightly different anecdote. However, once we find a result repeatedly across a series of simulated experiments, we can have more confidence in that the observed pattern is truly a *generic pattern* and readily accessible by evolution. With or without and understanding of the mechanistic details, relatively simple multilevel models can capture the eco-evolutionary dynamics of microbes, allowing us to study what happens, what else emerges from these dynamics “for free”, and equally important: what needs further explanation?

### Model overview

Virtual Microbes metabolise, grow and divide on a spatial grid (Figure 1C). Here, we use two parallel 40×40 grids with wrapped boundary conditions. One grid contains the Virtual Microbes and empty grid-points, and the other describes the local environment in which the Virtual Microbes live. This environmental layer holds influxed metabolites, waste products of Virtual Microbes, and spilled metabolites from lysing cells (Figure 1B). In order to express proteins, grow, and maintain their cell size, Virtual Microbes must synthesize predefined metabolite(s), which we call building blocks. These building blocks are not directly provided, but must be synthesized by the Virtual Microbes by expressing the right proteins, allowing them to pump / convert metabolites into one another (Figure 1A). The expression of these proteins depends on genes on genomes that undergo a wide variety of possible mutations upon reproduction (Table 1). Genomes are circular lists of genes, each with their own unique properties (*e.g*. K_*m*_, V_*max*_ for enzymes, K_*ligand*_ and binding motif for TFs). The level of expression is unique for each gene, and is determined by its evolvable basal transcription rate and how this rate is modulated by transcription factors. When an enzyme or transporter gene is expressed, that specific reaction will take place within the cell that carries that gene. Note however that in the complete metabolic universe, many more possible reactions exist. The genome of an evolved Virtual Microbes will typically only use a subset of all the possible reactions. Proteins to catalyse new reactions and novel TFs can be discovered through rare events.

**Table 1.** Types of mutations and their probabilities in WT evolution and serial transfer protocol (STP)

| Mutation | Description | Prob (WT evolution) | Prob (STP) |
| --- | --- | --- | --- |
| Duplication | A stretch of 1 or more genes is duplicated in tandem | 0.005 | 0.0015 |
| Deletion | A stretch of 1 or more genes is deleted | 0.005 | 0.0015 |
| Inversion | A stretch of 1 or more genes is inverted in order | 0.005 | 0.0015 |
| Translocation | A stretch of 1 or more genes is moved to a random location | 0.005 | 0.0015 |
| (stretch length) | Geometrically distributed with $p = 0.3$ | - | - |
| Gene discovery | Per time-step probability of discovering a new (randomly parameterised) gene. | 0.0002 | (disabled) |
| HGT | Per time-step probability of copying a gene from a cell closeby | 0.002 | (disabled) |
| Point mutation | Per gene per generation probability of modifying a single parameter of a gene (promoter strength, Michaelis Menten constants) | 0.005 | 0.0015 |
| Regulatory mutation | Per gene per generation probability of (partially) modifying the upstream binary operator sequence of a gene | 0.005 | 0.0015 |

Note that unlike most evolutionary models, fitness is not explicitly defined. Both the rate of birth and death are dynamically defined, and evolvable properties for Virtual Microbes. Birth depends on the availability of empty space and resources to synthesize building blocks, whereas death depends on the ability to survive under a variety of different conditions and the potential accumulation (and avoidance) of toxicity. The resulting survival of the fittest (referred to as “competitive fitness” by Fragata *et al.*, 2018) is an emergent phenomenon of eco-evolutionary dynamics[41].

#### Metabolic universe

The metabolic universe in Virtual Microbes is an automatically generated (or user defined) set of metabolites and reactions between them. The simple metabolic universe used in this study was automatically generated by a simple algorithm that defines 4 classes of molecules, how then can be converted into one another by means of 6 reactions, how fast they degrade, diffuse over the membranes, *etc*. (see Table 4).

The metabolism is simulated on the grid in terms of Ordinary Differential Equations (ODEs) using the Gnu Scientific Library in Cython. These ODEs include the influx of molecules into the system, diffusion between grid points, transport or diffusion across the membrane, intracellular metabolism (including expression and decay of proteins), biomass production, cell volume, *etc*.. Due to computational efficiency, the number of simulations was limited to 16 WTs and 16×3 “lab” experiments.

#### Transmembrane transport

For all molecules, transporters exist that import or export molecules across the cell membrane. Michaelis-Menten kinetics determine the transmembrane transportation with rate *v*:

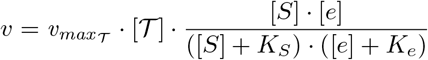

where 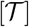 is the concentration of the transporter protein, [*S*] is the concentration of substrate transported, and [*e*] is the concentration of available energy carrier metabolites. *K_S_* and *K_E_* are the Michaelis-Menten constants for the substrate and energy carrier respectfully. Depending on the direction of transport (importing or exporting) [*S*] is either the external or the internal concentration of the substrate. Note that for any gene on the genome of a Virtual Microbe, 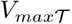, *K_S_* and *K_E_* are all freely evolvable parameters.

#### Metabolism

Similar to the transport, metabolic rates are catalysed by proteins by Michaelis-Menten kinetics with rate *v*:

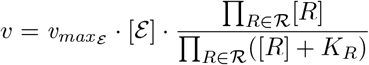

where 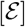 is the concentration of the enzyme catalysing the reaction, 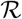 the set of all reactant metabolites, and *K_R_* and *v_maxε_* are evolvable kinetic parameters of enzyme 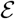.

#### Biomass production

Virtual microbes convert building block *B* to a biomass product *P*, which is consumed for cell growth and maintenance *Growth*(*B*) and protein production *Prod*(*B*), and determines strength with which individuals compete to reproduce. Biomass is next converted to cell volume with a fixed rate, and used for protein expression depending on the demands by the evolved genome. In other words, high rates of expression demand more biomass product for proteins, leaving less biomass product to invest in cell volume or maintainance (see cell volume growth). In total, the rate of change of *P* then becomes

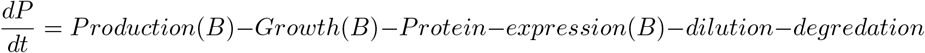

where B is the concentration of building block metabolites. Production is a lineair conversion of B into P, whereas growth, protein expression, and dilution depend on the dynamics of the cell. Biomass product is then consumed by cellular growth and protein expression which are a function of the building block concentration, is diluted proportional to the changes in cell volume, and degradation is fixed.

Consumption for protein expression is summed over all genes:

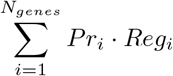

where *Pr_i_* is the basal expression rate of gene *i*, either up or down-regulated if transcription factors are bound to its operator sequence *Reg_i_* (see transcriptional regulation).

#### Cell volume growth

We assume that cell volumes a maximum cell size *MaxV* and that there is a continuous turnover *d* of the cell volume at steady state, ensuring the necessity to keep on metabolising even if there is no possibility to reproduce (*i.e.* if the grid points are all full). Volume then changes as

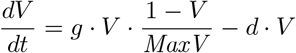

#### Transcriptional regulation

The rates at which genes are expressed is a function of the basal expression rate of the gene and the concentrations of binding TFs and their molecular ligands. The intrinsic basal expression rate of a gene is encoded by a strength parameter in a gene’s promoter region. This basal expression rate can be modulated by TFs that bind to an operator sequence associated with the gene. Binding sites and TF binding motifs are modelled as bit-strings and matching depends on a certain fraction of sequence complementarity. If a minimum complementarity is chosen < 1 a match may occur anywhere within the full length of the operator binding sequence and the TF binding motif. The maximum fraction of complementarity achieved between matching sequences linearly scales the strength with which a TF binds the target gene. In addition to binding strength following from sequence complementarity, TFs encode an intrinsic binding affinity for promoters *K_b_*, representing the structural stability of the TF-DNA binding complex.

TFs can, themselves, be bound to small ligand molecules with binding afinity *K_l_*, altering the regulatory effect they exert on downstream genes. These effects are encoded by parameters *eff_bound_* and *eff_apo_* for the ligand-bound and ligand-free state of the TF, respectively, and evolve independently. Ligand binding to TFs is assumed to be a fast process, relative to enzymatic and transcription-translation dynamics, and modeled at quasi steady state. We determine the fraction of TF that is not bound by any of its ligands *L*:

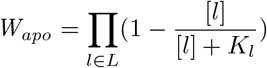

The fraction of time that a TF *τ* in a particular state *σ* (bound or apo) is bound to a particular operator *o*:

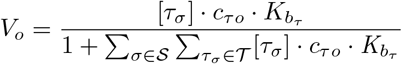

depends on the inherent binding affinity *K_bτ_* as well as the sequence complementarity score *c_τo_* between the tf binding motif and the operator sequence [cite Neyfahk]. The binding polynomial in the denominator is the partition function of all TFs 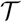 in any of the states 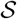 that can bind the operator. Note that small declines in the concentration of free TFs due to binding to operators are neglected.

Now, the operator mediated regulation function for any gene is given by

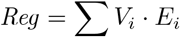

with *V_i_* the fraction of time that the operator is either unbound or bound by a TF in either ligand bound or unbound state and *E_i_* the regulatory effect of that state (1 if unbound or *eff_bound_* or *eff_apo_* when bound by a ligand bound or ligand free TF, respectively). Finally, protein concentrations 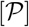 are governed by the function:

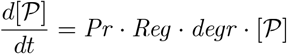

where *Pr* is the evolvable parameter *promoter strength* and *degr* a fixed protein degradation rate which is not evolvable.

#### Toxicity and death

Virtual Microbe death is a stochastic process depending on a basal death rate, which is potentially increased when internal metabolite concentrations reach a toxic threshold. A cumulative toxic effect is computed over the current life time *τ* of a microbe as

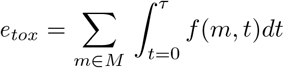

for all internal molecules *M*, with

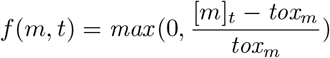

the toxic effect function for the concentration of molecule *m* at time *t* with toxicity threshold *tox_m_*. This toxic effect increases the death rate *d* of microbes starting at the intrinsic death rate *r*

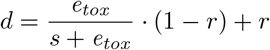

where *s* scales the toxic effect. Virtual Microbes that survive after an update cycle retain the toxic level they accumulated so far. Apart from toxicity and stochastic death, cells can also starve. When insufficient biomass product is availble to keep up the slowly decaying volume of the cell, the cells decrease in volume. If the cell volume drops below a *minimally viable volume*, this cell is automatically for death.

#### Reproduction

When cells compete for reproduction, the cells are ranked according to cell size. The “winner” is then drawn from a roulette wheel with weights proportional to this ranking. Upon reproduction, cell volume is divided equally between parent and offspring, and the genome is copied with mutations (see below). Molecule and protein concentrations remaining constant. Toxic effects built up during the parent’s lifetime do not carry over to offspring.

#### Genome and mutations

The genome is a circular list of explicit genes and their promoter region, organised like “pearl on a string”. Genes can be enzymes, transporters, or transcription factors. At birth, the genome is subject to various types of mutations. Large mutations include duplications, deletions, inversions, and translocations of stretches of genes (see Table 1). At the single gene level, point mutations allow all evolvable parameters to mutate individually (see Table 2). Horizontal gene transfer can occur on every time step. Innovations are an abstraction of “HGT from an external (off-grid) source”, and allow randomly parametrised genes to be discovered at any given moment with a low probability.

**Table 2.** Gene level mutations and the boundary conditions

| Parameter | Gene Types | Value range in simulation |
| --- | --- | --- |
| Promoter Strength | Enzyme, Transporter, TF | [0.001, 10] |
| $K_{substrate}$ | Enzyme, Transporter | [0.001, 10] |
| $K_{energy}$ | Transporter | [0.001, 10] |
| $K_{ligand}$ | TF | [0.001, 10] |
| $K_{operator}$ | TF | [0.001, 10] |
| $V_{max}$ | Enzyme, Transporter | [0.001, 10] |
| effect-bound | TF | [0.001, 10] |
| effect-apo | TF | [0.001, 10] |
| ligand | TF | A, B, C, or e |
| exporting | Transporter | True, False |
| sense-external | TF | [True, False] |
| binding-motif | TF | bit flip at random position |
| operator-sequence | Enzyme, Transporter, TF | bit flip at random position |

Note that unlike most evolutionary models, fitness is not explicitly defined. Both the rate of birth and death are dynamically defined, and evolvable properties for Virtual Microbes. Birth depends on the availability of empty space and resources to synthesize building blocks, whereas death depends on the ability to survive under a variety of different conditions and the potential accumulation (and avoidance) of toxicity. The resulting survival of the fittest (referred to as “competitive fitness” by Fragata *et al.*, 2018) is an emergent phenomenon of eco-evolutionary dynamics[41].

### Experimental setup

#### Metabolic network and wild type evolution

We use a very simple metabolic network with 2 resource metabolites, 1 building block metabolite, and an energy carrier (Figure 2A). We initialised 16 minimally viable Virtual Microbes, and evolved them for ~10.000-15.000 generations in fluctuating resource conditions by applying random fluctuations of the influx rates for the A and the C resource. Because the rate of influx for the two resource metabolites fluctuates between very high (10^−1^) and very low values (10^−5^), conditions can be very poor, very rich, and/or potentially toxic. To avoid total extinction, we subdivided the 40×40 grid into four 20×20 sub-spaces, in which these fluctuations are independent (see Figure 2B). Note however that these subspaces do not impede diffusion and reproduction, but merely define the rate at which resources flux into different positions on the grid. In this study, the microbes do not migrate during their lifetime. These conditions, summarized in Table 3, aim to simulate natural resource fluctuations, evolving what we call “wild types” (WTs) of Virtual Microbes. (see supplement S1)

**Table 3.** Grid setup and environmental forcing in WT evolution and serial transfer protocol (STP)

| Option (WT evolution) | Description | Value or range |
| --- | --- | --- |
| Maximum population size | As defined by the size of the grid (40x40) | 4900 |
| Sub-grids | The grid is sub-divided into n grids where fluctuations are independent | 4 |
| Fluctuation frequency | Probability (per time step) of 1 metabolite (A or C) changes in influx in one of the sub-grids | 0.01 |
| Fluctuation range | New influx of metabolite is sampled uniformly from range | [10e-5, 10e-1] |
| Extracellular metabolite outflux | Rate at which metabolites outside of cells wash out | 0.01 |
| Option (serial transfer protocol) | Description | Value |
| Maximum population size | As defined by the size of the grid (70x70) | 4900 |
| Number of cells serially transferred | A (near) tenfold dilution of cells | 500 |
| Time steps of cycle | This represents, for example, the “24 hour” serial transfer protocol of the LTEE | 50 (AUT) |
| [A] at beginning of cycle | Amount of resource A given at the beginning of the cycle | 1.25 |
| [C] at beginning of cycle | Amount of resource C given at the beginning of the cycle | 1.25 |
| Extracellular metabolite outflux | Assuming metabolites can no longer wash out of the system | 0.0 |

**Table 4.** A priori defined metabolites and reactions in artificial chemistry

| Metabolite | Mass | Class | Degradation rate | Diffusion rate | Toxicity level |
| --- | --- | --- | --- | --- | --- |
| A | 4 | Resource | 0.01 | 0.02 | 0.2 |
| B | 5 | Building block | 0.1 | 0.0015 | 0.2 |
| C | 6 | Resource | 0.01 | 0.015 | 0.2 |
| e | 1 | Energy carrier | 0.5 | 0.0015 | 0.2 |
| Potential reactions (6) |  |  |  |  |  |
| 1C → 1B + 1e | 1C → 1A + 2e | 1A + 1B → 1C | 2A → 1C | 2A → 1B | 1B → 1A + 1D |

The initial population consists of cells that have 3 enzymes, 3 pumps, and 5 transcription factors. All these proteins are randomly parameterized, meaning that these proteins are unlikely to have good binding affinities and catalytic rates. The amount of building block required to grow and produce protein is therefor very minimal in the early stages of evolution, and increases up to a fixed level as the Virtual Microbes improve.

#### In silico serial transfer protocol

We mimic a serial transfer protocol like that of the LTEE by taking our evolved WTs and – instead of fluctuating the resource conditions – periodically supplying a strong pulse of both the A- and the C-resource. While WTs are evolved in a spatial setting where resources flux in and out of the system, we here mix all cells and resources continuously and fully close the system, meaning no metabolites wash out. To apply strong bottlenecks while at the same time allowing for sufficient growth, we increased the size of the grid from 40×40 to 70×70. We dilute the approximately tenfold, transferring 500 cells to the next cycle. Firstly, horizontal gene transfer was disabled to represent the modified (asexual) *Escherichia coli* Bc251 clone that is used in the LTEE [1]. Furthermore, as the strong bottlenecks cause more genetic drift than the WT evolution, we found it necessary to dial back the mutation rates for the evolution of WTs to 30% to avoid over-exploiting mutants from appearing (see Table 1). Other parameters of the serial transfer protocol are listed in Table 3.

#### Growth rate and yield measurements

Yield was approximated by taking sum of all cell volumes, normalized by the maximum number of cells (i.o.w. the size of the grid). We measured yield both within a single serial transfer cycle (“daily yield”), and as the extended yield when we tested for long-term survival. As all WTs had slightly different log phases, we estimated the growth rates as the average building block production during the first half of the protocol.

#### Curating (quasi-)stable coexistence

Using the neutral lineage markers, we manually characterized coexistence by looking at the dynamics of neutral lineage markers. When two neutral markers had relatively stable frequencies with a daily pattern as visualised in Figure 5C for at least 10.000 time steps (approximately 100 generations), it was scored as coexistence. When this persisted for a while, but later got lost, it was scored as quasi-stable. When two neutral markers had balanced frequencies for at least 10.000 time steps and this pattern lasted until the end of the 800 serial transfers, it was scored as stable. If neither happened, it was annotated at no coexistence.

#### Further configuration of Virtual Microbes

Apart from the parameters within the confines of this article (Table 1-4), we have used the default settings for Virtual Microbes release 0.1.4, with the configuration files provided in Supplementary Section 2. Further details on the model and parametrisation are available online https://bitbucket.org/thocu/virtual-microbes

## Declarations

### Ethics approval and consent to participate

Not applicable

### Consent to publish

Not applicable

### Author contributions

B.D. performed simulations and provided the data. Results were analysed and interpreted by all authors. B.D. wrote the manuscript with input from P.H., J.M., and T.C.. P.H. supervised the project. All authors have approved the manuscript.

### Availability of data and materials

The full python module of Virtual Microbes is publicly available via PyPi. The code is available online on bitbucket.org/thocu/virtual-microbes. Further help with installation, instructions on how to use Virtual Microbes, and full documentation of the methods, is available on www.virtualmicrobes.com. As the data to support this study is fully computer generated, and consists of quite a large set of files, we felt it unnecessary and unhelpful to make the data available online. However, all the data that support this study are reproduced using the code and configuration from the supplementary materials. Finally, the corresponding author is available for help with the software.

## Acknowledgements

The authors want to thank Dominique Schneider and Thomas Hindré (Université Grenoble Alpes) for experiments done, and discussions had, during this project. Finally, the authors would like to thank Guillaume Beslon, and all the partners of the EvoEvo project for fruitful discussions.

## Supplementary materials - S1: Evolution of Virtual Microbes wild types

### Convergent and divergent evolution in Virtual Microbe wild types

In the evolution of our WTs we observed strong convergence as well as divergence in the metabolic and gene regulatory networks that evolved. Because the evolved populations consist of a rich mix of different genotypes, we here describe the WTs by profiling the gene repertoires and GRNs at the end of the simulation (~10.000 generations). For this, we took 20 (maximally unrelated) individuals from the evolved populations and determined the consensus metabolism (Figure S1A). While there is some diversity in the metabolic networks across WTs, the shared gene repertoire constitutes a metabolic network that forms a metabolic cycle complemented with resource importers and an exporter for the C metabolite (Figure S1B). We observed that the discovery of both the metabolic cycle as well as the exporter favour survival, as it coincides with an increase in population size and a decrease in the number of generations per time step (Figure S4). Note that in Virtual Microbes survival is improved by avoiding toxic effects of high metabolite concentrations and by only investing in growth when conditions are favourable for growth. The latter can be done via gene regulatory networks that respond to the quality of the environment, but we also found forms of metabolic regulation where microbes accurately fine-tuned kinetic parameters to automatically maintain homeostasis.

Although the shared gene repertoire from Figure S1B does not contain transcription factors (TFs), all of the 16 WTs have at least one type of TF. These TFs can constitutively repress or activate certain genes, or can respond to environmental conditions by binding to a ligand molecule. The latter response depends on the kinetic properties of the TFs and the properties of the genes which they regulate, all of which are evolvable (see Table S1). To get a better overview of how the WTs respond to environmental stimuli we therefore chose to directly measure the gene expression levels in a variety of different resource concentrations (displayed for 6 WTs in Figure S2). On the level of these GRNs, and their sensitivity the environment, we clearly see signs of strong divergent evolution. Note however, that the *effect* on the importer and exporter proteins seems very similar between WTs with different networks, showing that similar responses can be encoded by different GRNs. Finally, as seen in these graphs, some WTs have no response to environmental stimuli. We found that these non-regulating WTs are equally “fit”, in that they have the same rates of building block production and death rates (see Figure S3). However, the majority (11/16) WTs evolved clear regulatory mechanisms.

In short, during the *de novo* evolution of Virtual Microbe WTs, some evolved features seem highly predictable. Namely, all have evolved the metabolic cycle, all express both resource importer proteins, and all but one WT have a C-exporter. On the other hand, regulatory mechanisms and some of the secondary reactions display considerable diversity. Note that this divergence cannot be explained by differences in initial conditions or fluctuations in resource concentrations, because the WTs only differ with respect to the mutations that have happened in their evolutionary history. However, as shown in the main text, these differences have a profound effect on further evolution.

**Table S1.**
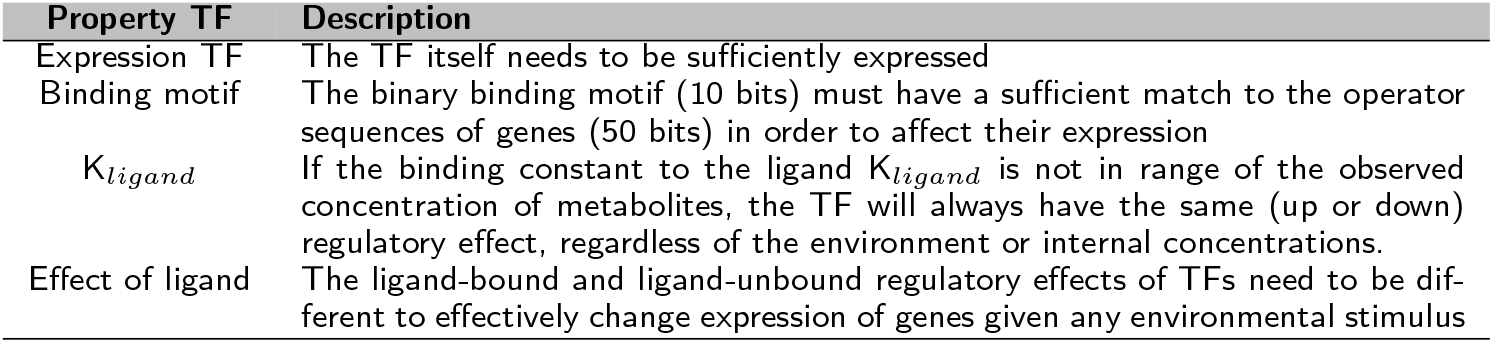
Important parameters for TFs for an environmental response.

**Table S2.** Anticipation and polymorphisms are also observed when changing in the serial transfer protocol. For different transfer times, dilutions, and resources concentrations, seven WTs (11-16, and WT07 from Figure 6 from the main text) have been tested for the anticipation effect and polymorphisms. Note that anticipation is not tested by prolonging the cycle (like in the main text), but by comparing the patterns in cell cycle dynamics with those from Figure 3 in the main text. If a clear decrease in cell volume was observed at the end of the cycle, it was scored as anticipation. Polymorphisms are scored as defined in the methods.

| Shorter transfer time (25), 800 cycles |  |  |
| --- | --- | --- |
| WT | Anticipation<br>(Yes/No) | Polymorph<br>(Yes, No, Quasi) |
| 11 | Y | N |
| 12 | Y | N |
| 13 | Y | Q |
| 14 | Y | N |
| 15 | Y | Q |
| 16 | Y | Q |
| 07 | Y | N |

| Higher transfer time (75), ~250 cycles |  |  |
| --- | --- | --- |
| WT | Anticipation<br>(Yes/No) | Polymorph<br>(Yes, No, Quasi) |
| 11 | Y | N |
| 12 | Y | N |
| 13 | Y | Y |
| 14 | Y | N |
| 15 | Y | N |
| 16 | Y | Q |
| 07 | Y | N |

## Supplementary materials - S2: Virtual Microbe configuration

The evolution of these WTs was done with Virtual Microbes version 0.1.4 which is publicly available as a Python package. Complete documentation on the methods is publicly available on http://bitbucket.org/thocu/virtualmicrobes. For this study, we used the configuration below. We removed options that are not relevant for reproducability (*e.g.* memory-limit, thread-count, data-storage-frequency etc.) or are default (*e.g.* universal-mutation-rate scaling=1.0) To reproduce these results with the newer versions of Virtual Microbes (0.2.4 as of January 2019), feel free to contact the corresponding author if help is required.

~~~
virtualmicrobes.py @reggen.cfg - evo @reg-evo.cfg --name My_WT_Vmicrobe
~~~

genera_options.cfg:

~~~
– – base – death – rate 0.01
– – mutation – rates
chrom_dup = 0.0
chrom_del = 0.0
chrom_fiss = 0.0
chrom_fuse = 0.0
point_mutation = 0.005
tandemdup = 0.005
stretch_del = 0.005
stretch-invert = 0.005
stretch_translocate = 0.005
stretch_exp_lambda = 0.3
external_hgt = 0.0002
internal_hgt = 0.002
regulatory_mutation = 0.005
reg_stretc_ex_lambda = 0.1
– – point – mutation – ratios
ligand_class = 0.1
exporting = 0.1
– – rand – gene – params
base = 10
lower = – 1.0
upper = 1.0
– – mutation – param – space
base = 10
lower = –0.5
upper = 0.5
min = 0.01
max = 10.
– – max – historic – max 0.1
– – growth – rate – scaling 1
– – competition – scaling 1
– – selection – pressure historic_window_scaled
– – historic – production – window 1000
– – scale – prod – hist – to – pop
– – small – mol – diff – const 0.02
– – prot – degr – const 0.7
– – –– transporter – membrane – occupancy .1
– – influx – range
base = 10
lower = –1.0
upper = –5.0
– – fluctuate – frequencies 0, 0.01
– – init – external – conc 0.
– – small – mol – ext – degr – const 1e – 2
– – bb – ext – degr – const 1e – 1
– – ene – ext – degr – const 5e – 1
– – –– transcription – cost 0.002
– – energy – transcription – scaling 0.01
– – spill – conc – factor 1.
– – v – max – growth 1
– – min – bind – score 0.85
– – per – grid – cell – volume 8
– – enzyme – volume – occupancy 3
– – grid – sub – div row = 2, col = 2
– – sub – env – part – influx 1.0
– – gri d – cols 40
– – grid – rows 40
~~~

evosim_options.cfg:

~~~
– – duration 1000000
– – env – rand – seed 87
– – reproduce – size – proportional
– – cell – init – volume 1.5
– – cell – growth – rate 2.0
– – cell – shrink – rate 0.6
– – cell – growth – cost 0.2
– – cell – division – volume 2.
– – init – prot – mol – conc .1
– – max – cell – volume 5
– – nr – resource – classes 3
– – nr – energy – classes 1
– – ene – energy – range 1 , 1
– – res – energy – range 2 , 10
– – nr – building – blocks 1
– – building – block – stois 1 , 1
– – nr – cell – building – blocks 1
– – mol – per – ene – class 1
– – mol – per – res – class 1
– – nr – cat – reactions 3
– – nr – ana – reactions 3
– – max – nr – cat – products 2
– – min – cat – energy 1 , 3
– – max – nr – ana – products 1
– – nr – ana – reactants 2
– – chromosome – compositions tf = 0, enz=1, pump= 1
– – binding – seq – len 10
– – operator – seq – len 50
– – prioritize – influxed – metabolism
– – init – prot – mol – conc 0.01
– – degradation – variance – shape 100
– – no – toxicity – variance – shape
– – toxicity 0.2
– – toxic – building – blocks
– – toxicity – scaling 1000
– – tf – binding – cooperativity 2
– – homeostatic – bb – scaling 1
– – high – energy – bbs
– – prioritize – energy – rich – influx
~~~

**Figure S1.**
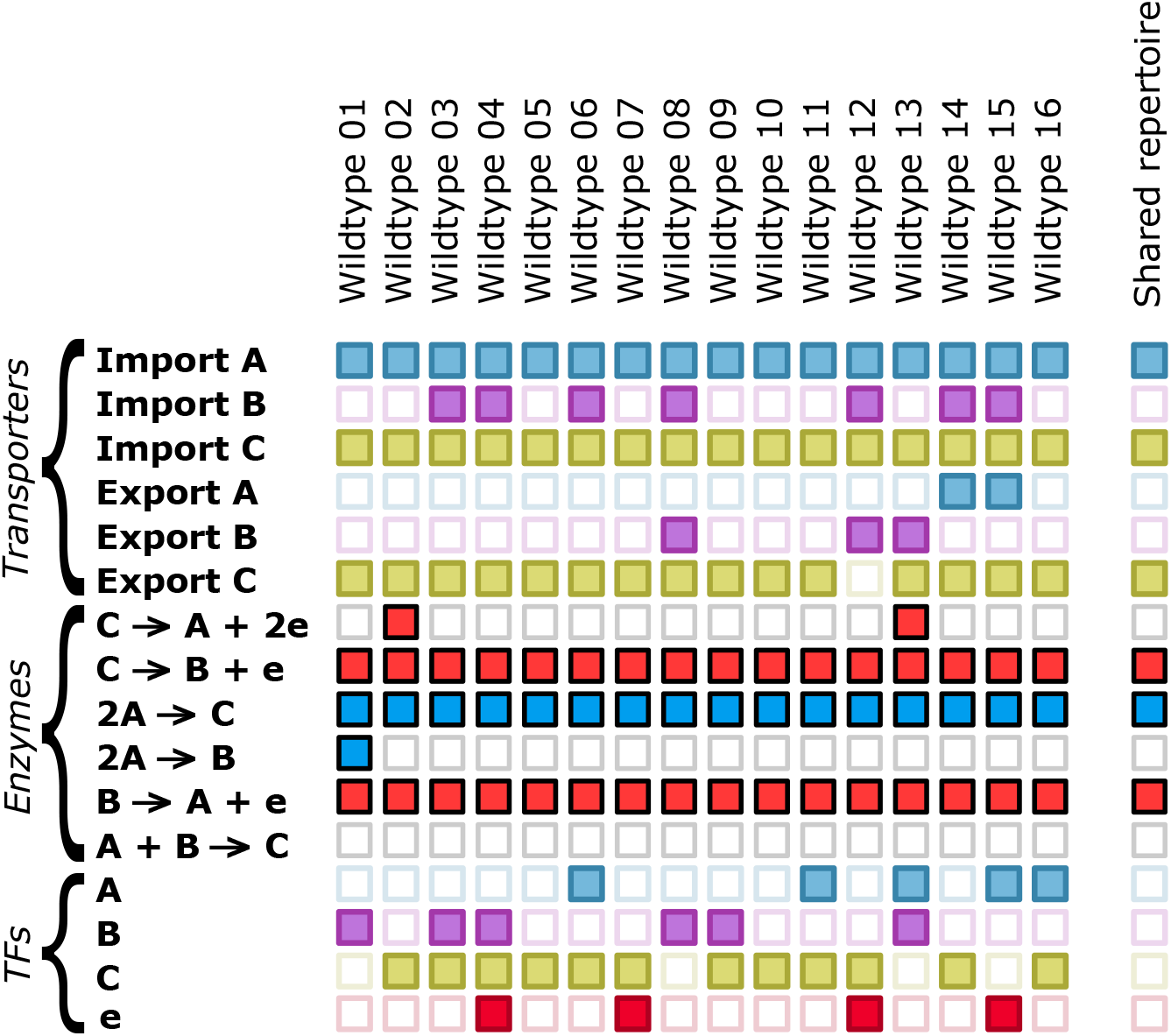
The evolved gene repertoires for all 16 WTs. The gene repertoires of WTs (20 maximally unrelated individuals) is displayed for all 16 replicate simulations after 1.000.000 time steps. Rows represent the different types of proteins (transporters, enzymes and TFs), and the columns the gene repertoires. Note that the presence of a gene does not imply it is functional, since properties such as K_*s*_ and V_*m*_*ax* might be poorly parameterised. Genes found in less than 4 cells or genes with low concentrations (*i.e.* low expression) were omitted. The seperate column depicts the genes found in at least 90% of WTs.

**Figure S2.**
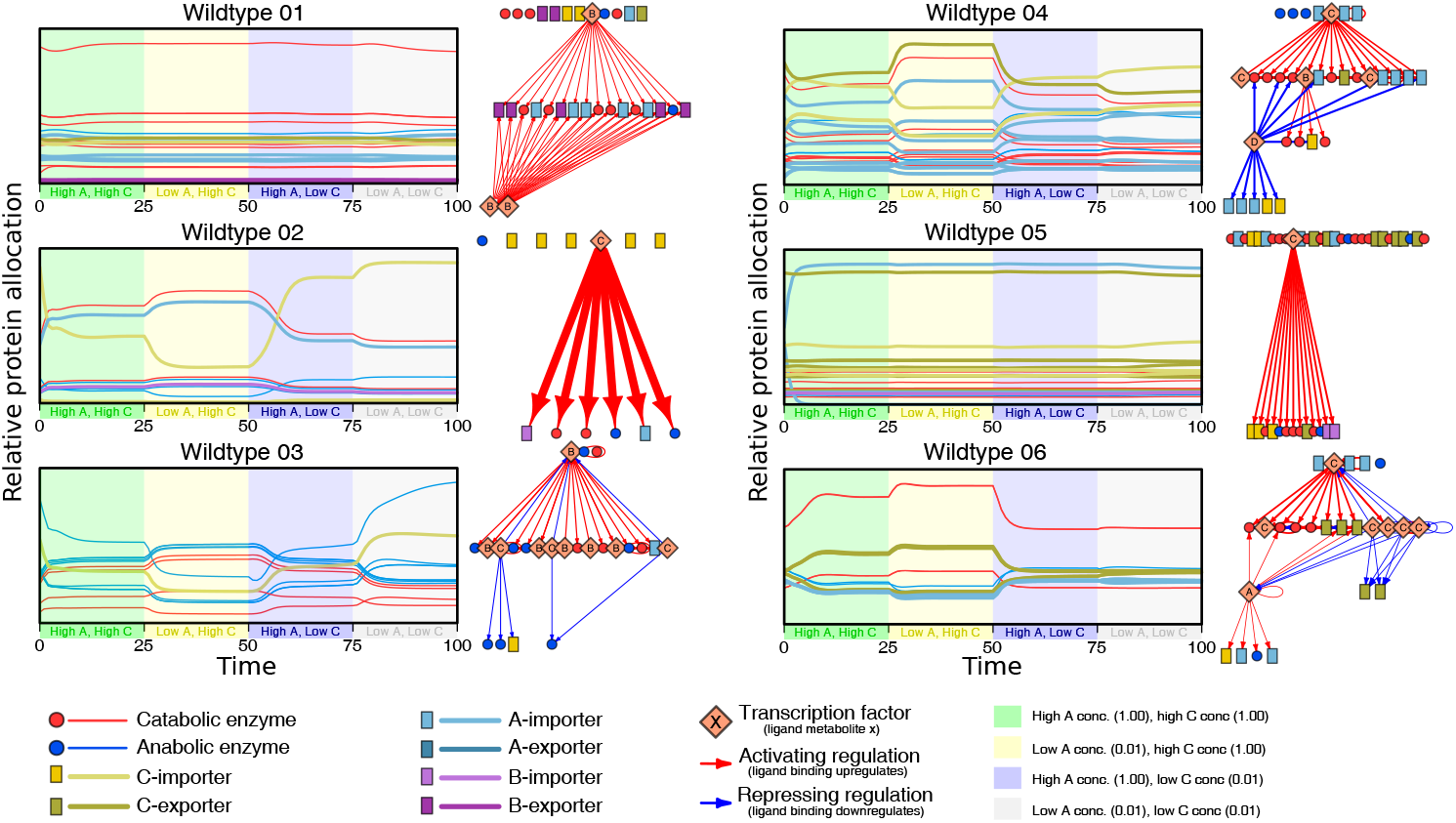
WTs have great diversity of transcriptional regulation, and not all respond to changing resource concentrations. The graphs shows how the protein allocation shift to imposed changes in resource concentration of the environment. Single clones from the WTs were taken and resource concentrations of the A- and C-resource were varied from low (0.01) to high (1.0). The gene regulatory network responsible for these changes is displayed next to each graph. The colours for different enzymes are as displayed in the legend. Thicker arrows in the gene regulatory network represent higher expression levels of the transcription factors. Genes with very low expression levels were omitted for clarity.

**Figure S3.**
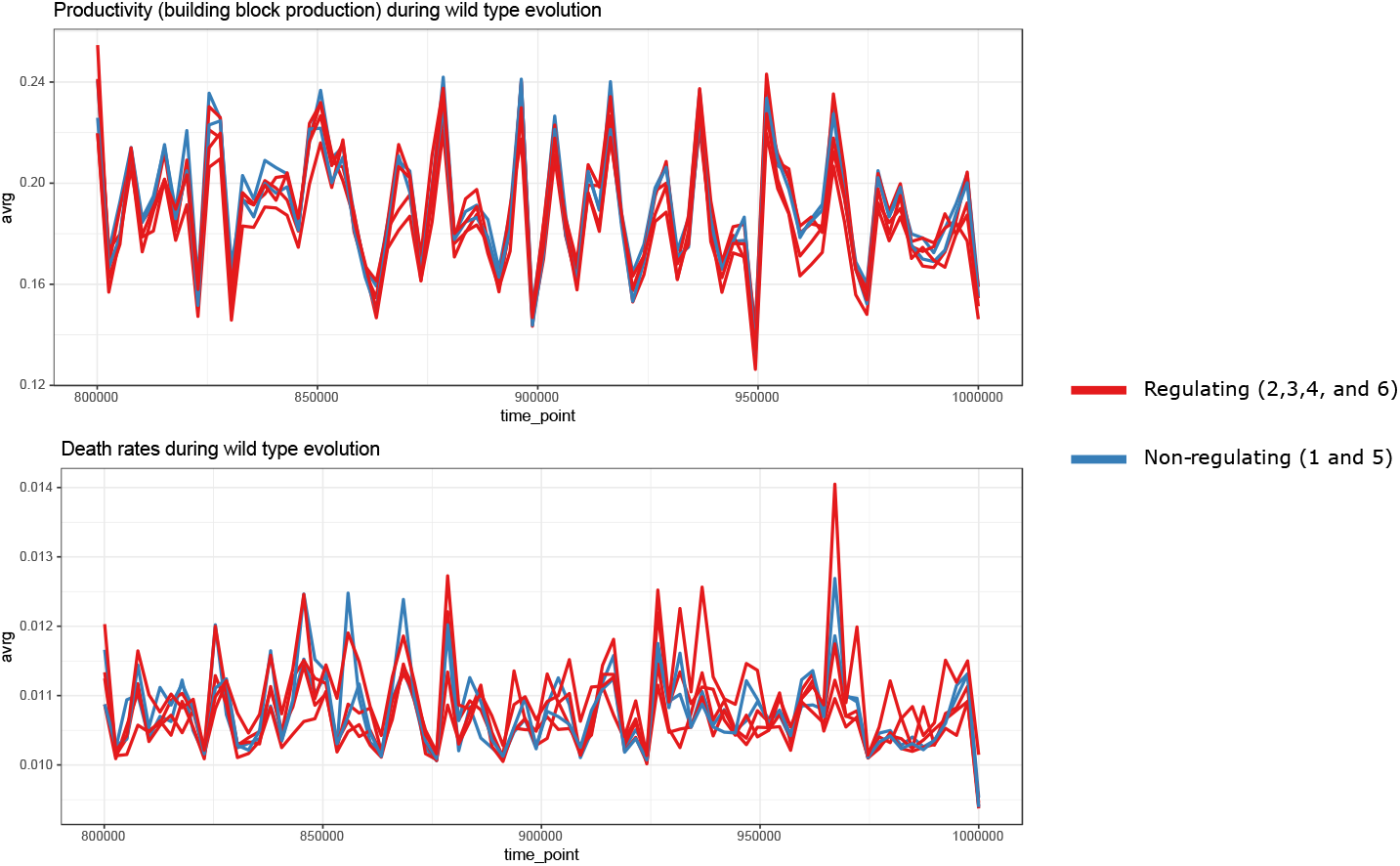
Different WTs have similar building block production rates and death rates. For 6 WTs, the last 2000 generation of WT evolution did not show any differences in “fitness”, which is approximated by the rates of building block production and death. Furthermore, regulating WTs showed not difference compared to non-regulating WTs.

**Figure S4.**
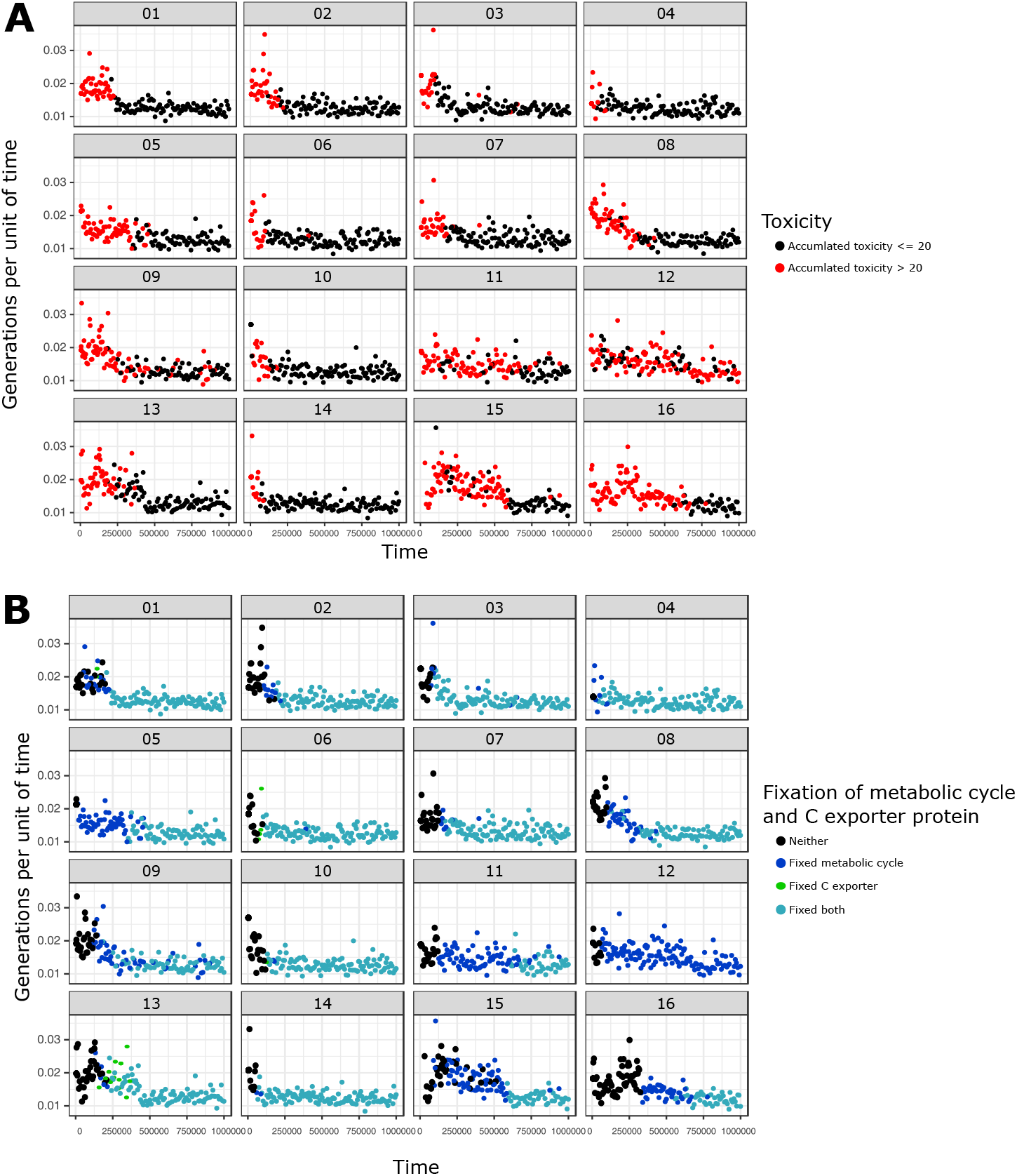
Decrease in number of generations per time step for WTs coincides with a decrease in toxicity and/or fixation of a metabolic cycle. Decrease in number of generations per time step coincides with a decrease in toxicity accumulation and/or fixation of a metabolic cycle. Every dot represents an average over a 100 generations of simulation. For panel A, red dots represent a toxicity level above which death rate is increased at least threefold (toxicity > 20), and black dots represent the lower toxic levels (toxicity < = 20). In panel B, the cyan dots represent the fixation of both the C-exporter and the metabolic cycle, green dots the fixation of only the C-exporter, blue dots the fixation of the metabolic cycle, and black dots the fixation of neither of these features.

**Figure S5.**
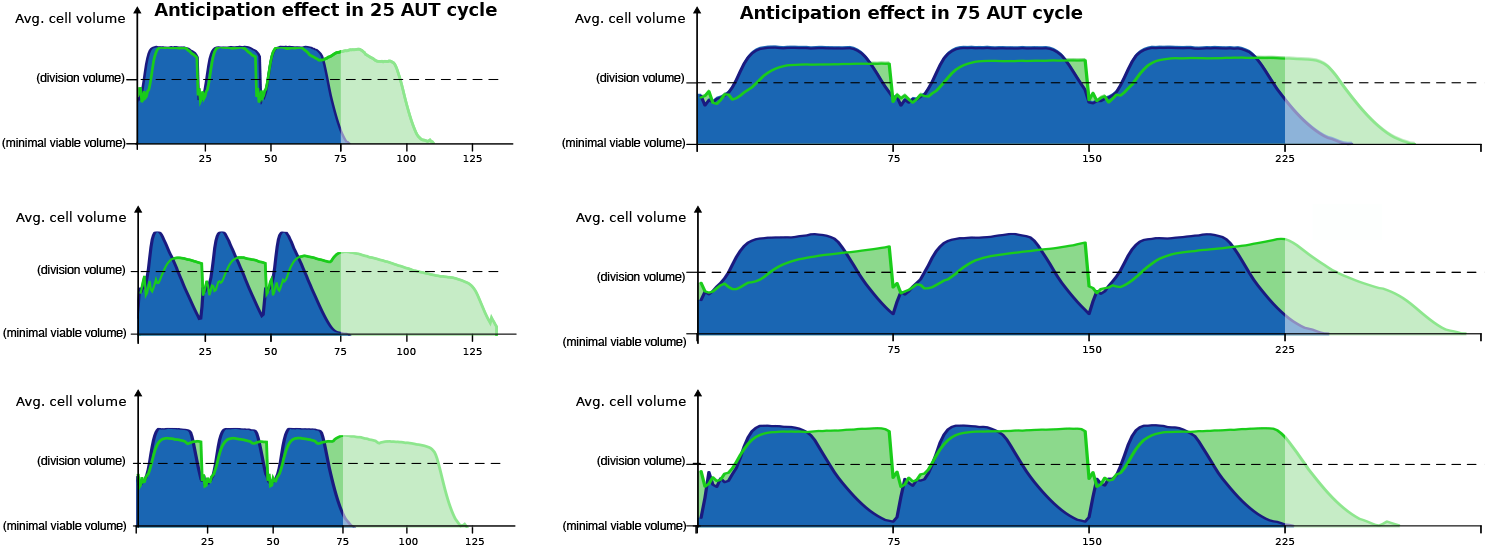
The anticipation effect is present after evolving in shorter (left) and longer (right) time intervals.

**Figure S6.**
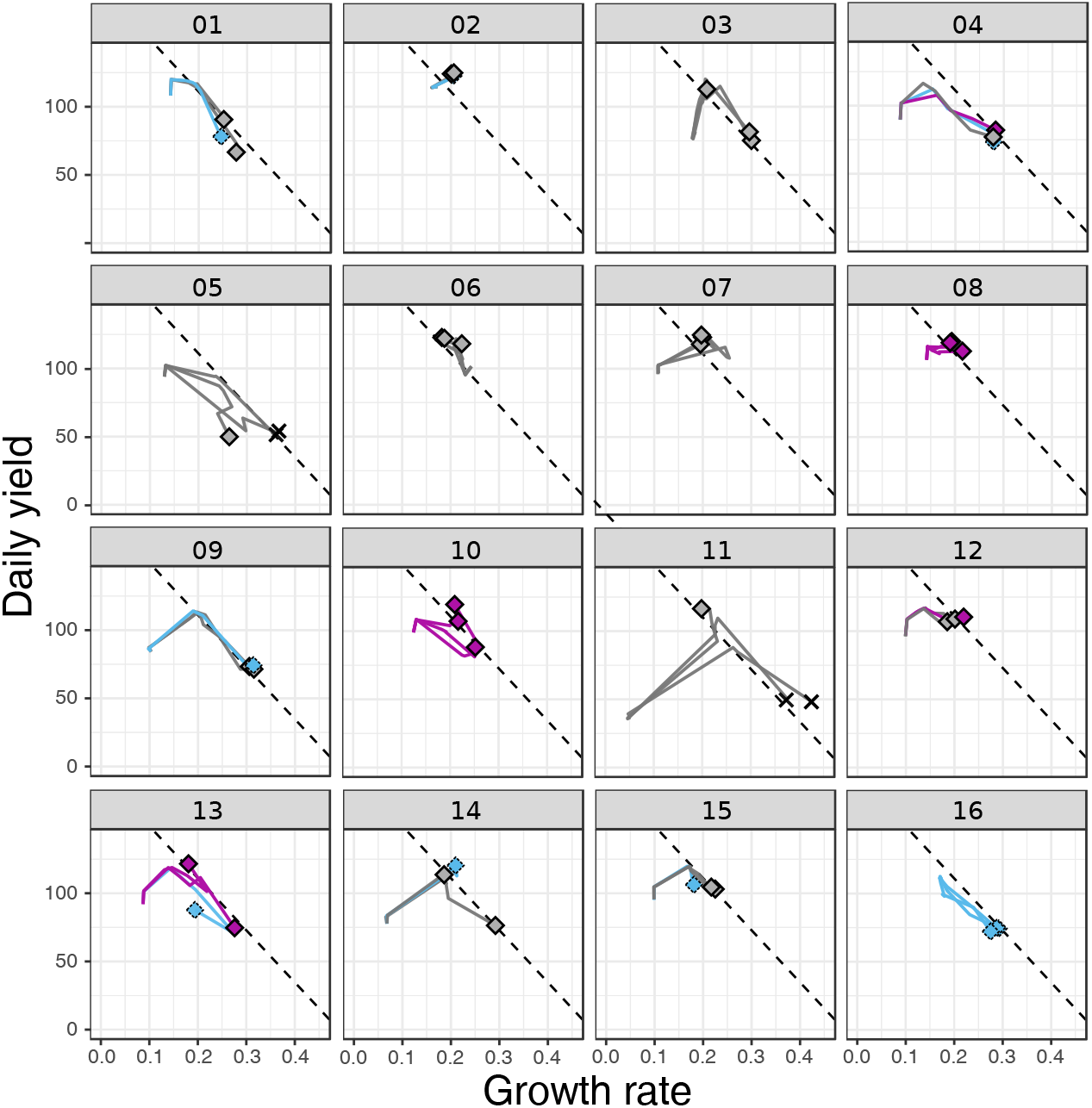
Three replicate serial transfer experiments for all 16 WTs. This image is shows all the data from the examples from Figure 4.

**Figure S7.**
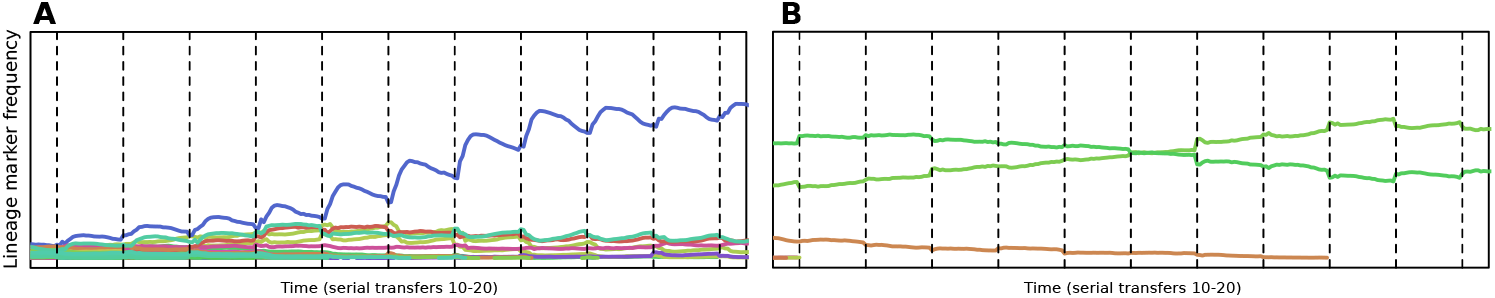
Dynamics of different invading mutants. A) Neutral lineage markers linked to mutations which cause high growth rates but poor survival increase in frequency early, but are losing in frequency during stationary phase. B) Neutral lineage markers linked to mutations that have higher growth rates without trading off against survival show no such temporal fitness effect.

**Figure S8.**
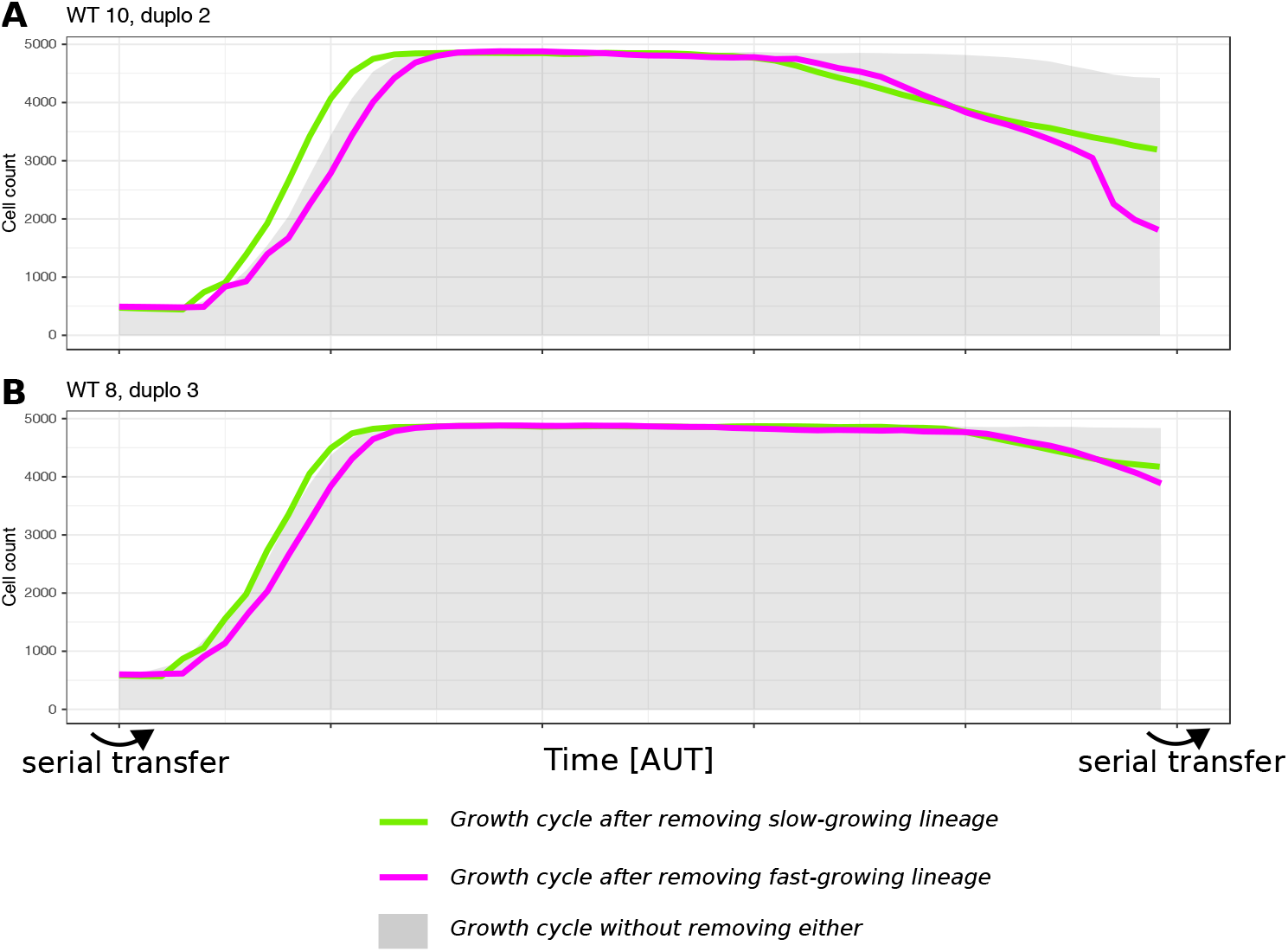
Lineages based on resource specialisation (top) and growth vs. yield specialisation (bottom) perform better in the presence of the other. In the upper panel, two resource specialists over-exploit their resource in the absence of the other. In the bottom panel, without evident resource specialisation, both lineages have increased toxicity (given that this happens through enzymatic changes, this is likely due to changes in fluxes), which accumulates to significant levels near the end of the cycle. This figure thus shows that lineages can grow dependent on one another, as they are part of each-others environment.

